# Polysialic Acid-Functionalized MAP Scaffolds Promote Regulatory Immune Responses After Ischemic Stroke

**DOI:** 10.1101/2025.09.10.674054

**Authors:** Yunxin Ouyang, Sanchuan Che, Emma C. Whitehead, Koravit Poysungnoen, Anshu Agarwal, Albert Liu, Hannah Newman, Peter Silinski, Matthew L. Becker, Tatiana Segura

## Abstract

Glycosylation regulates immune and neural functions within the central nervous system (CNS), yet biomaterials rarely leverage glycans due to their structural complexity. Polysialic acid (PSA), comprising α2,8-linked sialic acid residues, is a promising candidate owing to its potent immunomodulatory interactions with inhibitory Siglec receptors. Systematic screening of multiple sialic acid derivatives identifies PSA as uniquely effective in inducing anti-inflammatory polarization of bone marrow-derived macrophages (BMDMs). Based on these findings, an injectable microporous annealed particle (MAP) scaffold presenting PSA covalently via its reducing end (MAP-PSA) is engineered, recapitulating physiological glycan orientation. MAP-PSA exhibits robust mechanical properties, stable glycan immobilization, and resistance to enzymatic degradation. Using ischemic stroke as a CNS injury model, MAP-PSA significantly reduces neutrophil infiltration and inflammatory activation while enhancing reparative macrophage and microglial phenotypes. These immunomodulatory effects persist into subacute stages, characterized by sustained reductions in inflammation and enhanced microglial homeostasis. Overall, MAP-PSA scaffolds demonstrate a novel therapeutic paradigm for CNS injuries such as stroke, with translational potential for broader neuroinflammatory and regenerative applications.

## 1. Introduction

Glycosylation, the enzymatic attachment of glycans to proteins or lipids, is essential for regulating immune and neural signaling within the CNS ^[1]^. Despite their central biological importance, glycans remain underutilized in biomaterials, which predominantly rely on delivering proteins, cells, or small molecules. These conventional approaches frequently face limitations, including limited stability, complex formulation requirements, challenging delivery routes, and stringent storage conditions, hindering clinical translation^[2]^.

Glycan-based biomaterial approaches offer compelling advantages, as glycans offer lower susceptibility to degradation during storage compared to proteins and modulate immune responses through receptor interactions. Crucially, as natural post-translational modifications, glycans simultaneously modulate multiple pathways^[1b]^. This multifunctionality contrasts with protein- or small molecule-based approaches, which often require intricate combinations and precise dosing of multiple factors to achieve equivalent biological outcomes^[3]^. Thus, integrating bioactive glycans into biomaterial scaffolds can yield simplified therapeutic platforms, thereby reducing formulation complexity.

Extensive studies have highlighted polysialic acid (PSA), a polymer of α2,8-linked Neu5Ac, as a crucial regulator of immune responses in the CNS^[4]^. Previous biomaterial strategies incorporating PSA primarily employed nanoparticles or non-specific conjugation methods, achieving immune modulation and regeneration mainly in peripheral models such as spinal cord injury or ocular disease^[5]^. However, these approaches have not utilized biomaterial platforms designed for sustained local retention, precise receptor engagement, or direct immune-cell interactions within CNS lesions. Specifically, for ischemic stroke, existing biomaterial approaches have predominantly used soluble Neu5Ac monomers as drug carriers rather than as direct immunomodulatory signals^[6]^. Physiologically, PSA is covalently attached at its reducing end to cell-surface proteins, orienting the non-reducing end outward to interact efficiently with immune receptors such as inhibitory Siglecs (sialic acid binding Ig-like lectins)^[7]^.

To better leverage PSA’s immunoregulatory potential, we engineered an injectable microporous annealed particle (MAP) hydrogel scaffold composed of hyaluronic acid (HA), conjugating PSA specifically via its reducing end, closely mimicking the physiological orientation of PSA found naturally on cell surfaces^[7]^. The MAP scaffold’s porous architecture enables controlled spatial delivery directly to CNS lesion sites such as ischemic stroke infarcts, facilitating immune cell infiltration and sustained modulation of immune responses at clinically relevant therapeutic windows^[8]^. This biomimetic design aims to enhance targeted engagement of inhibitory Siglec receptors expressed by infiltrating immune cells, resulting in potent and sustained immunomodulation favorable for CNS regeneration^[4a]^.

In this study, we first systematically evaluated the immunomodulatory potential of PSA relative to other sialic acid derivatives using bone marrow-derived macrophages (BMDMs). Subsequently, PSA-functionalized MAP scaffolds demonstrated robust control over inflammatory responses in an ischemic stroke mouse model, significantly reducing pro-inflammatory neutrophil infiltration and activation, while enhancing reparative macrophage and microglial phenotypes. Further immunohistochemical analyses at a later subacute stage post-stroke confirmed sustained spatial modulation of immune cell infiltration, highlighting reduced inflammation and enhanced microglial responses. Overall, this study presents a biomimetic glycan-based biomaterial approach for ischemic stroke therapy, demonstrating significant improvements over prior glycan biomaterial strategies and highlighting broad translational potential for CNS regeneration.

## 2. Results and Discussion

### 2.1. Sialylation as a Dynamic Regulator of CNS Immunity and Injury Responses

Glycans are increasingly appreciated as dynamic regulators of cellular processes, significantly influencing cellular signaling ^[9]^. Major glycan classes include N-glycans (linked via asparagine residues) and O-glycans (linked via serine or threonine residues), each exhibiting distinct patterns of sialylation—addition of terminal sialic acids (Neu5Ac)—that influence cellular signaling and immune interactions (Figure 1a)^[9a^, ^10^]. N-glycan sialylation varies significantly by species and brain region, whereas mucin-type O-glycans, found on proteins like Podoplanin and MUC1, exhibit consistently high sialylation (>90%), emphasizing their role in local neuroimmune interactions ^[11]^. These distinct sialylation patterns highlight complementary functions of N- and O-glycans in CNS homeostasis.

**Figure 1.**
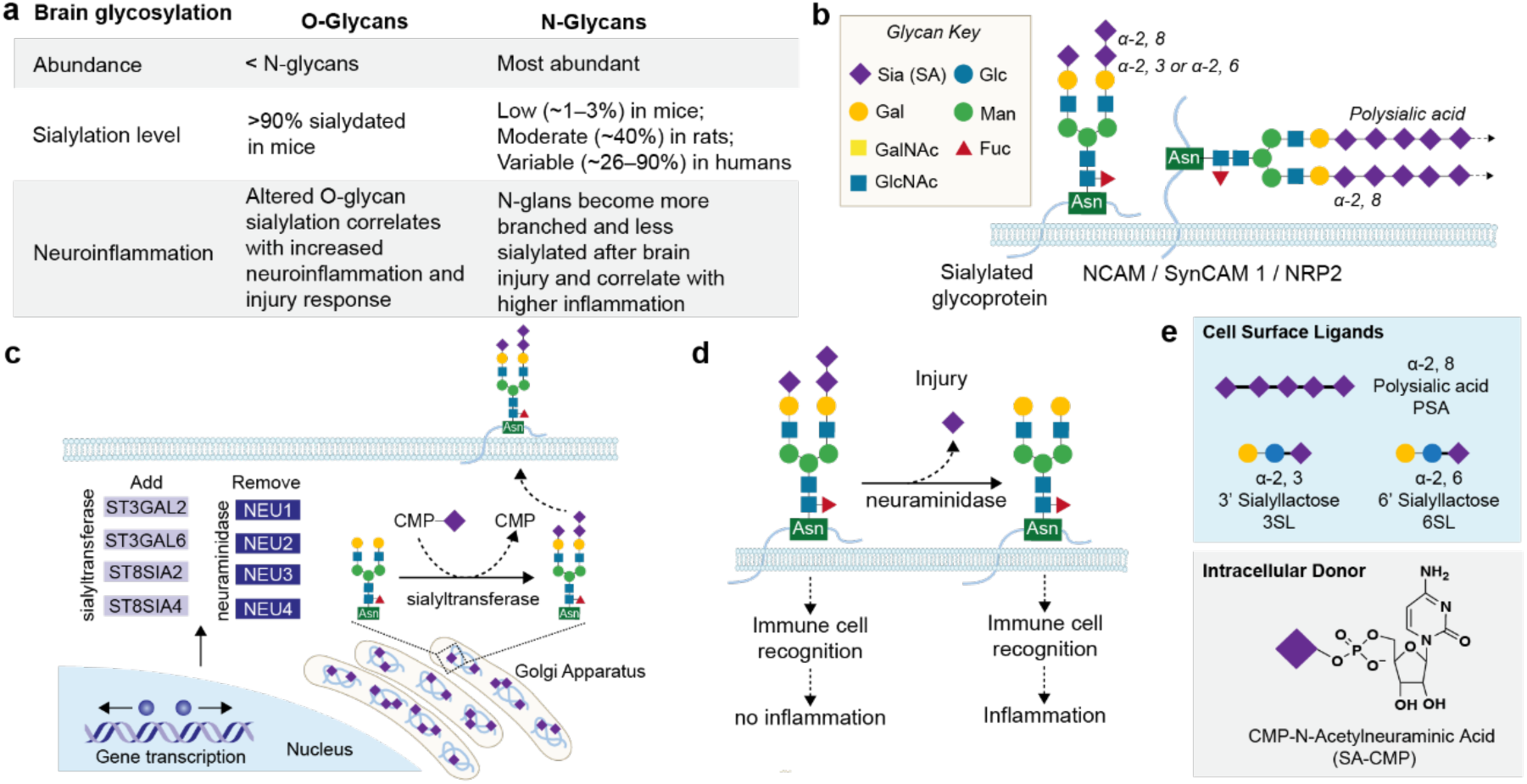
Sialylation landscape in the brain and its potential for immunomodulation. a) Comparative overview of brain glycan abundance and sialylation levels across species, highlighting marked differences between O- and N-glycans and variability across detection methods and species-specific contexts. b) Schematic of representative N-glycan structures, including sialylated and polysialylated (α2,8-linked) forms attached to neural glycoproteins such as NCAM, SynCAM 1, and NRP2. c) Biosynthesis pathway of sialylated glycans, emphasizing Golgi-localized enzymatic processes and subsequent trafficking to plasma membranes. d) Impact of neuraminidase-mediated de-sialylation after neural injury, resulting in altered immune cell recognition and enhanced inflammation. e) Chemical structures and functional categories of sialic acid derivatives utilized to investigate immune modulation in macrophages, distinguishing cell-surface ligands (PSA, 3’SL, 6’SL) from intracellular Golgi substrates (CMP-SA).

In the CNS, sialic acids predominantly form α2,3-, α2,6-, or α2,8-linkages (Figure 1b). Specifically, α2,3- and α2,6-linked sialic acids modify key glycoproteins involved in neuron–glia communication and inflammatory signaling, such as integrin β1 and CD22**^[12]^**. PSA uniquely modifies select CNS proteins, including neural cell adhesion molecule (NCAM), SynCAM 1, and neuropilin-2 (NRP2), critically regulating neurodevelopmental and neuroimmune responses**^[7]^**.

Sialylation of glycoproteins primarily occurs within the Golgi apparatus **via** sialyltransferases (e.g., ST3Gal2, ST3Gal6, ST8Sia2, ST8Sia4), utilizing cytidine monophosphate-Neu5Ac (CMP-Neu5Ac) as the sugar donor (Figure 1c). Newly sialylated glycoproteins are trafficked to the plasma membrane, where their terminal sialic acids directly engage inhibitory Siglec receptors on immune cells, regulating inflammatory signaling**^[13]^**. After CNS injuries like stroke, increased neuraminidase activity cleaves terminal sialic acids, reducing inhibitory Siglec engagement and intensifying inflammation, microglial/macrophage activation, and secondary tissue damage (Figure 1d)**^[13–14]^**.

Concurrently, the extracellular matrix (ECM) composition dramatically shifts, with elevated chondroitin sulfate proteoglycans (CSPGs) creating inhibitory environments for regeneration**^[15]^**. Restoring or preserving sialylation thus offers therapeutic potential to mitigate harmful inflammation and enhance CNS repair.

To systematically explore immunomodulation via sialylation, we selected sialic acid derivatives: PSA for its potent engagement with inhibitory Siglec^[16]^, 3’-sialyllactose (3’SL), and 6’-sialyllactose (6’SL) representing common shorter linkages^[17]^, and the intracellular donor CMP-Neu5Ac (CMP-SA)^[18]^, an intracellular donor to evaluate intracellular versus extracellular sialylation effects (Figure 1e).

### 2.2. Endotoxin Removal Enables Accurate Evaluation of PSA-mediated Immunomodulation

Commercial PSA, derived from Escherichia coli fermentation, contains endotoxin contaminants (Figure 2a), which activate inflammatory signaling through Toll-like receptor 4 (TLR4)^[13^, ^19]^. To remove endotoxins, we used a Triton X-114 phase separation method (Figure 2b)**^[20]^**. PSA Triton X-114 treatment (PSAx) significantly reduced endotoxin contamination in the PSA from >4000 EU/mg to approximately 1.34 EU/mg (Figure 2a)^[20]^, which is a value below the FDA recommended endotoxin value for brain implants**^[21]^**.

**Figure 2.**
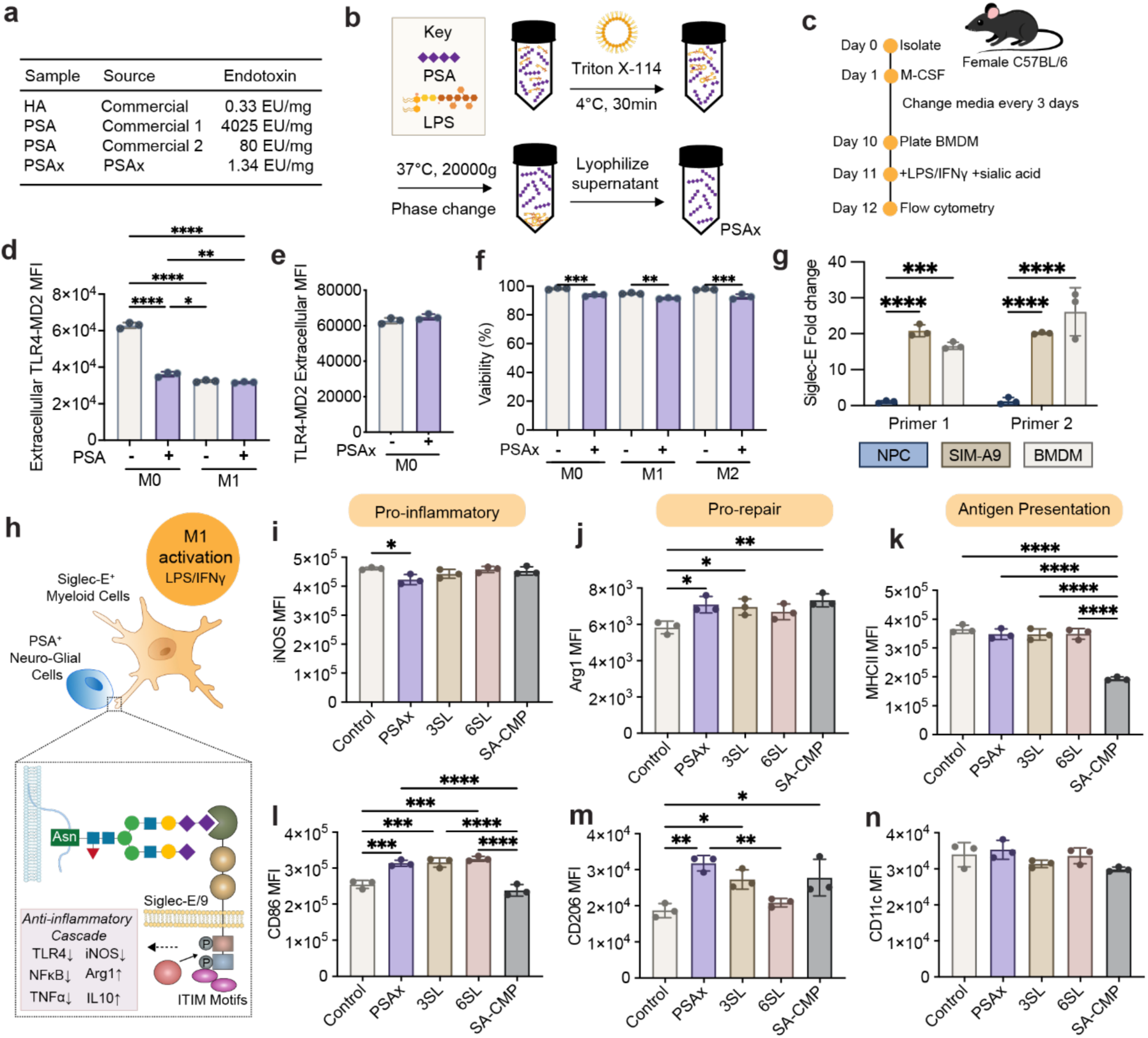
Endotoxin removal and screening of sialic acid derivatives for immunomodulatory activity. a) Endotoxin levels of commercially obtained polysialic acid (PSA) before and after purification via Triton X-114 phase separation (PSAx), compared to hyaluronic acid (HA). b) Schematic representation of endotoxin removal from commercial PSA. c) Experimental timeline for macrophage culture, M1 polarization (LPS + IFN-γ), and addition of sialic acid derivatives. d–e) Flow cytometric analysis of extracellular TLR4-MD2 expression on bone marrow-derived macrophages (BMDMs) with untreated PSA (d) and endotoxin-removed PSAx (e). f) BMDM viability assessed by flow cytometry under M0, M1, and M2 conditions with or without PSAx treatment. g) qPCR analysis of Siglec-E expression in BMDM, microglial SIM-A9 cells, and neural progenitor cells (NPCs) using two independent primer sets. h) Schematic illustrating PSA interactions with inhibitory Siglec receptors on myeloid cells, highlighting downstream anti-inflammatory signaling pathways. i–n) Flow cytometry quantification of macrophage polarization markers (iNOS, Arg1, MHCII, CD86, CD206, CD11c) under M1 activation with soluble sialic acid derivatives: PSA, 3’SL, 6’SL, and CMP- SA. Data presented as mean ± SEM, n = 3 biological replicates: *p<0.05, **p<0.01, ***p<0.001, ****p<0.0001, determined by one-way ANOVA with Tukey’s post-hoc test.

As expected, PSA, straight from the manufacturer, results in TLR4 activation (via flow cytometry) in bone marrow-derived macrophages (BMDMs) cultured under unstimulated (M0) conditions, while PSAx does not (Figure 2c). PSA significantly reduced extracellular TLR4-MD2 levels nearly two-fold, indicative of receptor internalization similar to LPS/IFN-γ-induced (M1) activation (Figure 2d)**^[22]^**. Endotoxin-removed PSAx restored extracellular TLR4-MD2 expression to baseline, confirming this receptor internalization was endotoxin-mediated rather than inherent to PSA itself (Figure 2e). Additionally, cell viability remained consistently high (>90%) despite minor reductions following purification, potentially due to residual detergent effects (Figure 2f). These data highlight the importance of endotoxin removal to reliably assess PSA-driven immunomodulation.

### 2.3. Systematic Screening of Sialic Acid Derivatives for Immunomodulatory Potential

To identify the most suitable sialic acid (SA) derivative for modulating inflammation, we screened four derivatives with known affinities for Siglec receptors: PSA, 3’SL, 6’SL, and the intracellular donor CMP-SA. Murine Siglec-E binds primarily α2,8-linked SA, while human Siglec-9 and Siglec-7 recognize α2,3/α2,6 and α2,8 linkages, respectively^[23]^.

To confirm our macrophage model expressed relevant receptors, we quantified Siglec-E expression in BMDMs relative to neural progenitor cells (NPCs) as a negative control and the microglial cell line SIM-A9 as a positive control (Figure 2g). qPCR analysis consistently showed significantly elevated Siglec-E mRNA in both BMDMs (∼16-fold increase) and SIM- A9 microglia (∼20-fold increase) compared to NPCs.

Interaction between Siglec-E on myeloid cells and PSA initiates inhibitory signaling via phosphorylation of immunoreceptor tyrosine-based inhibitory motifs (ITIMs), recruiting phosphatases SHP-1/SHP-2 to suppress pro-inflammatory pathways (NF-κB, iNOS) and enhance anti-inflammatory markers (Arg1, IL-10) (Figure 2h)^[23b, 24]^. To directly test whether soluble SA derivatives influence macrophage phenotype, we treated bone marrow-derived macrophages (BMDMs) under classical pro-inflammatory activation (M1: LPS + IFN-γ). PSA uniquely suppressed the pro-inflammatory marker iNOS versus activated control (Figure 2i), while simultaneously enhancing reparative markers Arg1 and CD206 (Figure 2j, 2k). PSA also moderately increased CD86 expression without significantly altering MHCII (Figure 2l, 2m). In contrast, 3’SL and 6’SL strongly elevated CD86 (∼2-fold), however their effects on Arg1 and CD206 are less pronounced than PSA. CMP-SA, an intracellular donor, robustly increased Arg1 and CD206 similarly to PSA, but distinctly decreased MHCII (Figure 2j–m). In the absence of stimulation (M0), Arg1 expression increased significantly with PSA, 6’SL, and CMP-SA, while CD86 expression notably increased with 6’SL and CMP-SA. Under M2 polarization, CMP-SA again markedly increased Arg1 and CD86 expression but significantly reduced CD11c, highlighting its robust intracellular modulation of glycosylation pathways.

Given these distinct immunomodulatory effects, PSA was selected for biomaterial integration based on its pronounced suppression of inflammatory markers (iNOS), enhanced reparative polarization (Arg1, CD206), and receptor-mediated mechanism (Siglec-E).

Practical advantages further supported PSA selection. Its polymeric structure tolerates partial cleavage during chemical modifications, maintaining sufficient chain length for receptor interactions**^[25]^**. Additionally, its higher molecular weight simplifies purification via dialysis compared to shorter glycans, which require more complex chromatography^[26]^. Intracellular CMP-SA relies heavily on glycosylation substrates (e.g., terminal galactose residues) and enzymes that are compromised during stroke-induced inflammation, limiting its therapeutic viability ^[27]^. Thus, we focused on developing a biomaterial platform that could achieve sustained, physiologically relevant PSA presentation, closely mimicking natural glycan- receptor interactions.

### 2.4. Site-specific Chemical Modification Enables Physiological Mimicry of PSA Presentation

Endogenous PSA presentation involves covalent attachment at the reducing end, orienting its non-reducing ends outward for receptor engagement^[7]^. To mimic this orientation, we developed a controlled, reducing-end-specific chemical modification strategy coupled with bio-orthogonal conjugation, enabling controlled PSA immobilization onto MAP scaffolds.

We initiated PSA modification through reductive amination at its reducing end (Figure 3a). PSA was first reacted with cysteamine dihydrochloride under mildly basic conditions (pH 8.4, 55 °C) to open the reducing-end hemiacetal ring, forming a Schiff base intermediate^[28]^. Subsequent reduction with sodium cyanoborohydride and thiol reduction using dithiothreitol yielded PSA-thiol (PSA-SH). We then selectively oxidized PSA’s non-reducing end, generating reactive aldehydes, and performed aniline-catalyzed oxime ligation with aminooxy-functionalized biotin, producing biotin-labeled PSA-SH^[29]^. Finally, bio- orthogonal inverse electron-demand Diels–Alder click chemistry between the thiol group on PSA-SH and a maleimide-PEG-tetrazine linker produced PSA-SH-tetrazine derivatives^[30]^. This modular chemistry route provided dual functional handles, allowing both conjugation and detection of PSA.

**Figure 3.**
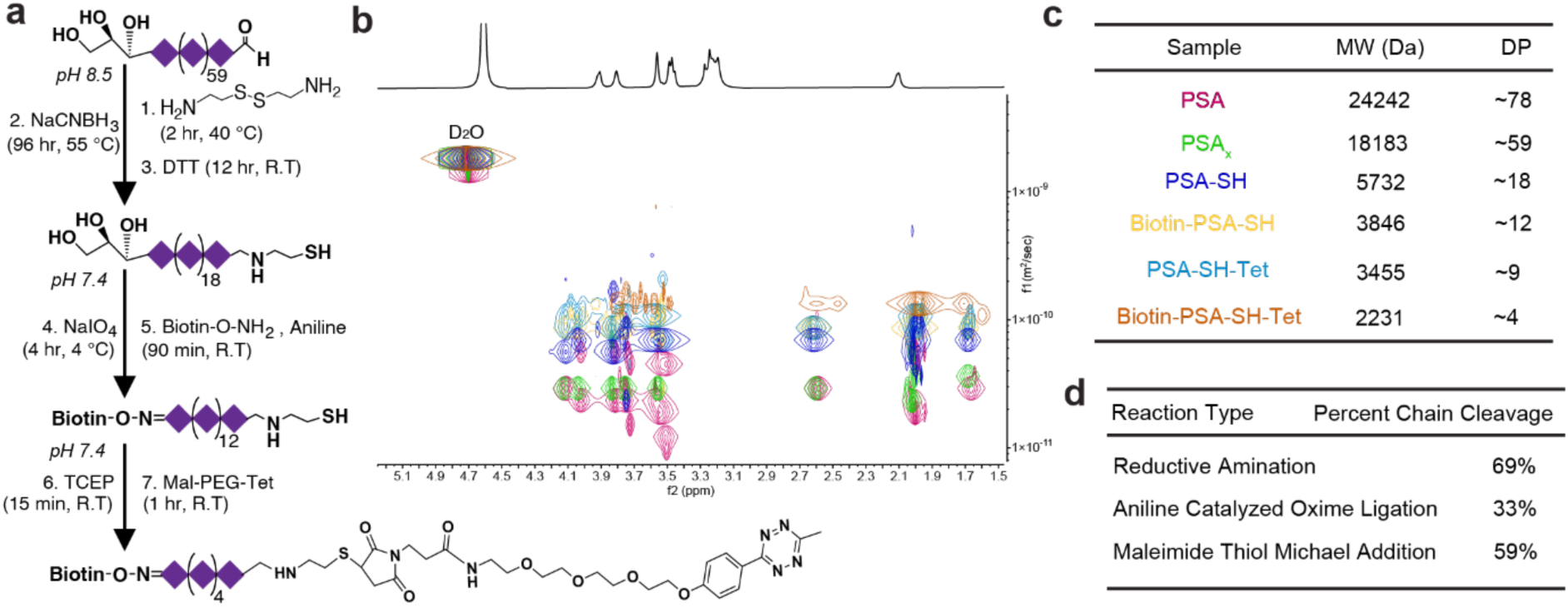
Site-specific chemical modifications generate PSA derivatives with controlled functionalization. a) Stepwise synthesis: reducing-end reductive amination, non-reducing end oxidation, and tetrazine addition. b–c) DOSY-based molecular weight and degree of polymerization characterization demonstrate progressive polymer shortening. d) Calculated chain cleavage percentages identify reaction-specific fragmentation.

The modification process resulted in progressive PSA chain cleavage, likely due to elevated temperature and prolonged incubation, conditions known to induce glycosidic bond hydrolysis. We used diffusion-ordered spectroscopy (DOSY) to characterize the degree of cleavage (Figure 3b), correlating measured diffusion coefficients with PEG (polyethylene glycol) standards (Figure S2). We observed progressive reduction in PSA molecular weight and degree of polymerization (DP) after each modification step. PSA molecular weight decreased from ∼24 kDa (DP_n_ = 78) to ∼5.7 kDa (DP_n_ = 18) after reductive amination, ∼3.8 kDa (DP_n_ = 12) after oxime ligation, and ∼2.2 kDa (DP_n_ = 4) after maleimide-thiol reaction (Figure 3c–d). Reductive amination induced the greatest cleavage (69%), while subsequent reactions caused additional, lesser fragmentation. Previous studies report varied functional PSA polymer lengths in physiological contexts, from short oligomers (∼2–7 residues) to long chains (∼50 residues) ^[31]^. Shorter PSA chains (DP_n_ ∼8–20) have shown effective immunomodulatory effects, such as regulating immune recognition and modulating inflammatory responses, relevant in contexts of innate immune activation ^[31b]^. Thus, the PSA derivatives generated here (DP_n_ ∼ 4 – 18) fall within previously reported biologically relevant ranges. Subsequent studies are needed to confirm whether these shortened PSA derivatives retain desired immunomodulatory activities when presented on scaffolds.

We next validated controlled immobilization of PSA onto hydrogel microparticles (HMPs). Spherical microparticles (∼80–90 µm diameter) were synthesized from norbornene-modified hyaluronic acid (HA-NB) as previously reported (Figure 4a)^[32]^. PSA-tetrazine derivatives were then conjugated to residual norbornene groups on HA-HMPs through bio- orthogonal inverse electron-demand Diels–Alder click chemistry, achieving PSA immobilization specifically via its reducing end (Figure 4b). Fluorescence imaging using Alexa Fluor 568-labeled PSA confirmed successful conjugation on the particle surfaces (Figure 4b).

**Figure 4.**
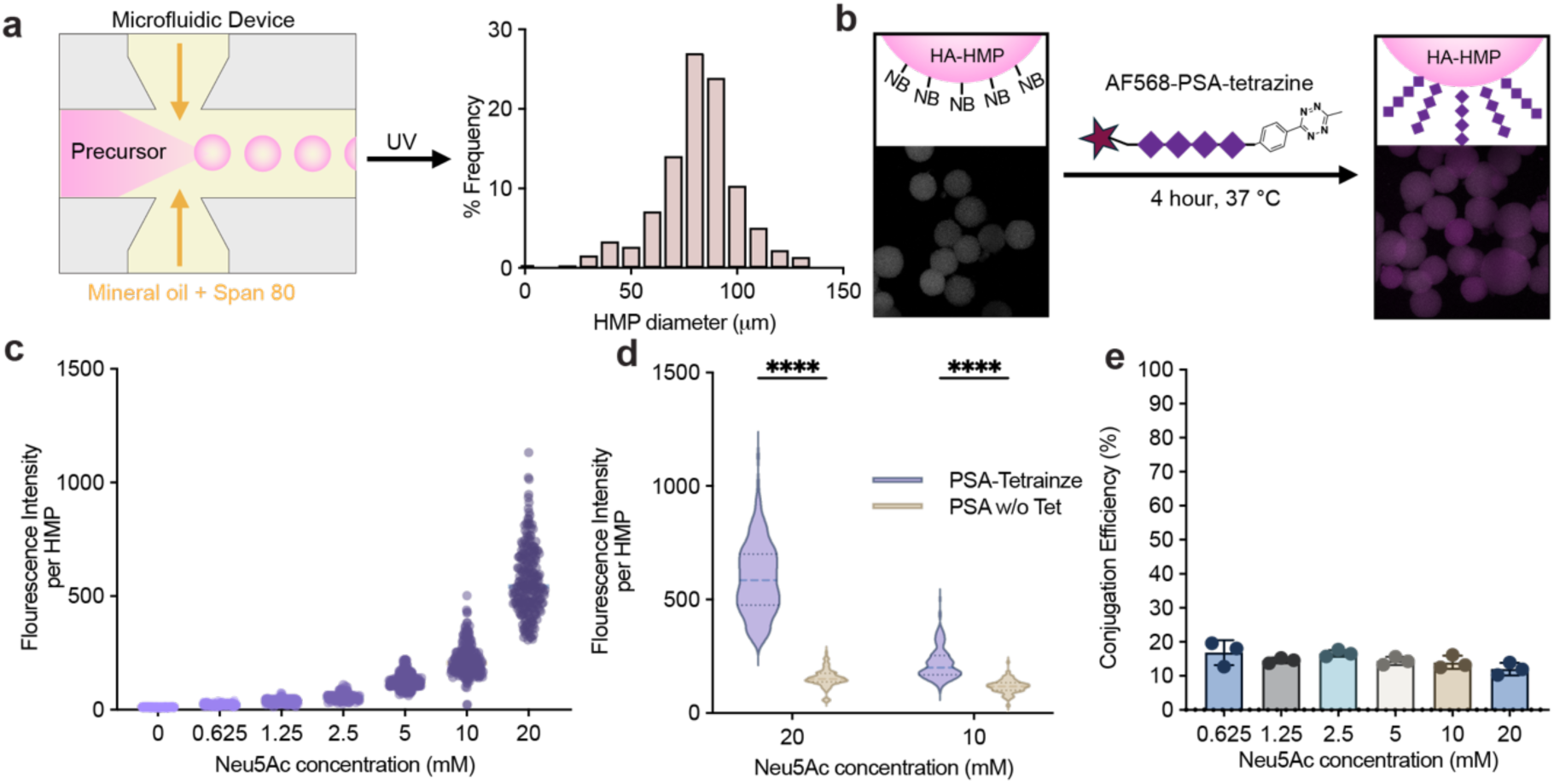
Bio-orthogonal conjugation enables tunable and stable PSA immobilization on hydrogel microparticles. a) Microfluidic preparation of HA-HMPs with consistent size distribution. b) Schematic and fluorescence images confirm successful PSA conjugation via tetrazine-norbornene chemistry. c–e) Quantification of conjugation reveals tunable PSA density (c), specificity imparted by tetrazine chemistry (d), and consistently high efficiency across various conditions (e). Data in (c, d) represent individual HMP measurements (n > 100 HMP per concentration, pooled from 3 independent functionalization batches). Data in (e) represent mean ± SEM from three independent functionalization batches; no significant differences observed by one-way ANOVA with Tukey’s post-hoc test.

Quantitative fluorescence analysis demonstrated a concentration-dependent increase in PSA immobilization (Figure 4c). PSA conjugation efficiency was significantly higher with tetrazine-modified PSA compared to PSA lacking tetrazine, confirming specificity provided by the bio-orthogonal approach (Figure 4d). Conjugation efficiency was consistent (∼13–17%) across PSA concentrations (Figure 4e). Slight reductions at high concentration may reflect diffusion limitations, though further reaction optimization could enhance conjugation efficiency. This bio-orthogonal approach enabled controlled PSA immobilization onto HMPs.

### 2.5. PSA-MAP Retains Mechanical Integrity and Enzymatic Stability

We next used rheology to assess how PSA conjugation affects scaffold mechanical integrity. MAP scaffolds were prepared by mixing PSA-HMP and unmodified HA-HMP (both types of HMPs were modified with Alexa Fluor 568–PSA-tetrazine) at specific ratios (0%, 50%, and 100% PSA-HMP), followed by annealing with HA-tetrazine (HA-Tet) to form microporous scaffolds, where HA-Tet covalently crosslinks residual norbornene groups on individual HMPs. Confocal imaging confirmed uniform distribution and spatial heterogeneity of PSA-HMPs within the scaffolds (Figure 5a).

**Figure 5.**
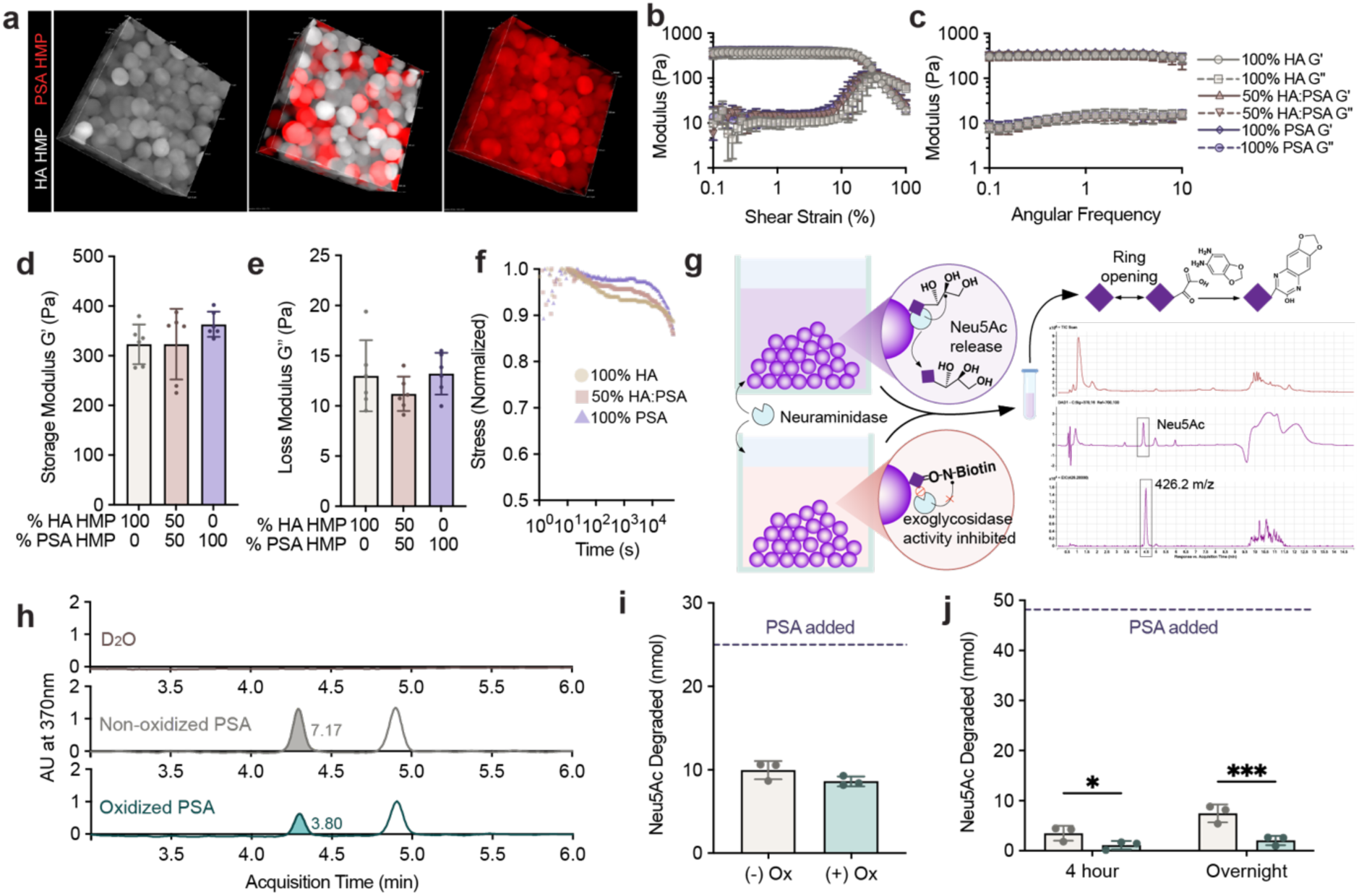
PSA conjugation maintains scaffold mechanical integrity and oxidation significantly improves enzymatic stability. a) Confocal imaging demonstrating tunable PSA distribution and scaffold homogeneity. b–c) Rheological characterization via amplitude and frequency sweeps confirm minimal mechanical differences upon PSA incorporation. d–e) Quantified storage (G’) and loss (G’’) moduli remain consistent across scaffold types, matching neural tissue stiffness. f) Stress-relaxation assays reveal subtle differences in scaffold relaxation profiles with increasing PSA content. g) Schematic depiction of sialidase-mediated PSA degradation assay and detection of released Neu5Ac by DMB derivatization and HPLC-MS. h) Representative chromatograms confirm reduced Neu5Ac release from oxidized PSA scaffolds. i–j) Quantitative analysis of Neu5Ac release after enzymatic degradation: (i) soluble, unconjugated PSA polymer incubated with sialidase, and (j) PSA conjugated to MAP scaffolds incubated with sialidase, demonstrating significantly enhanced enzymatic resistance conferred by non-reducing-end oxidation under physiologically relevant conditions. Data presented as mean ± SEM (n=3–5 independently casted scaffolds per formulation). One-way ANOVA with Tukey’s post hoc test for (d-e) and student’s t-test for (i-j): *p < 0.05, **p < 0.01, ***p < 0.001.

Rheological characterization revealed minimal impact of PSA conjugation on scaffold mechanical properties. Amplitude sweeps indicated similar linear viscoelastic ranges across scaffold compositions (Figure 5b). Frequency sweep assays showed consistent viscoelastic behavior across conditions (Figure 5c). Storage (G’) and loss (G’’) moduli remained statistically comparable among scaffold compositions (G’ ∼300–400 Pa), confirming appropriate scaffold stiffness mimicking brain tissue mechanics (Figure 5d-e)^[33]^. Stress-relaxation profiles exhibited subtle differences, with PSA-containing scaffolds showing slightly delayed relaxation, potentially reflecting changes in network interactions due to PSA conjugation (Figure 5f) ^[34]^.

We next used computational analysis (LOVAMAP software) to quantify pore architecture (Figure S4a)^[35]^. PSA incorporation did not significantly alter overall scaffold porosity, as demonstrated by comparable void volume fractions across formulations (Figure S4b). Additional pore metrics (pore volume, surface area, characteristic lengths, aspect ratio, average internal diameter, and longest pore length) remained similar across scaffolds (Figure S4c-h. Previous work indicates that pore size impacts macrophage phenotype^[36]^; thus, stable pore characteristics ensure the observed immunomodulatory effects result from PSA presentation rather than unintended structural differences.

Given PSA’s susceptibility to enzymatic degradation by neuraminidases, which are upregulated post-stroke^[14c]^, we assessed the stability of PSA incorporated into MAP scaffolds under enzymatic conditions. MAP scaffolds containing either oxidized or non-oxidized PSA were incubated with sialidase, followed by quantification of released free sialic acid (Neu5Ac) via derivatization and detection by HPLC-MS (Figure 5g). Representative chromatograms showed clear peaks corresponding to DMB-labeled Neu5Ac at approximately 4.3 min retention time. The peaks were visually lower in MAP conjugated with oxidized PSA compared to non-oxidized PSA, suggesting oxidation conferred protection from sialidases (Figure 5h).

At physiologically relevant enzyme concentrations (0.2 U/µL), neither oxidized nor non-oxidized PSA-conjugated scaffolds exhibited detectable degradation products after overnight incubation, indicating robust stability under typical cell culture conditions (Figure S3 d-e)^[37]^. At higher enzyme concentrations (4 U/µL), oxidized PSA consistently showed reduced degradation compared to non-oxidized PSA at both 4-hour and overnight time points (Figure S3 f-i). These observations were validated by quantitative analyses from integrated peak areas and standard curves. For soluble PSA polymers, oxidized PSA displayed slightly lower, though not statistically significant, degradation compared to non-oxidized PSA after overnight incubation (Figure 5i). However, measurements of MAP scaffold supernatants showed that oxidation significantly enhanced enzymatic stability, releasing approximately three-fold less Neu5Ac compared to non-oxidized PSA after both 4-hour and overnight incubations at higher enzyme concentrations (Figure 5j).

Together, these results underscore the utility of oxidation at the non-reducing end of PSA for enhancing scaffold resilience to sialidase-mediated degradation. Given that physiologically relevant sialidase concentrations did not degrade non-oxidized PSA- conjugated scaffolds, we proceeded with non-oxidized PSA scaffolds for subsequent studies^[37–38]^. This decision was also supported by the fact that prior work showing that non-oxidized PSA maintains a higher degree of polymerization, which is potentially beneficial for biological activity^[31b, 39]^.

### 2.6. PSA-MAP Modulates Macrophage Polarization Toward Regulatory Phenotypes *In Vitro*

We next encapsulated BMDMs within PSA-MAP scaffolds to test their efficacy in modulating macrophage polarization (Figure 6a). BMDMs were encapsulated within scaffolds containing varying PSA densities (0, 0.625, 2.5, and 20 mM Neu5Ac monomer equivalents) then stimulated with pro-inflammatory M1 cytokines (LPS and IFN-γ). Confocal microscopy confirmed homogeneous macrophage distribution within the scaffolds (Figure 6b).

**Figure 6.**
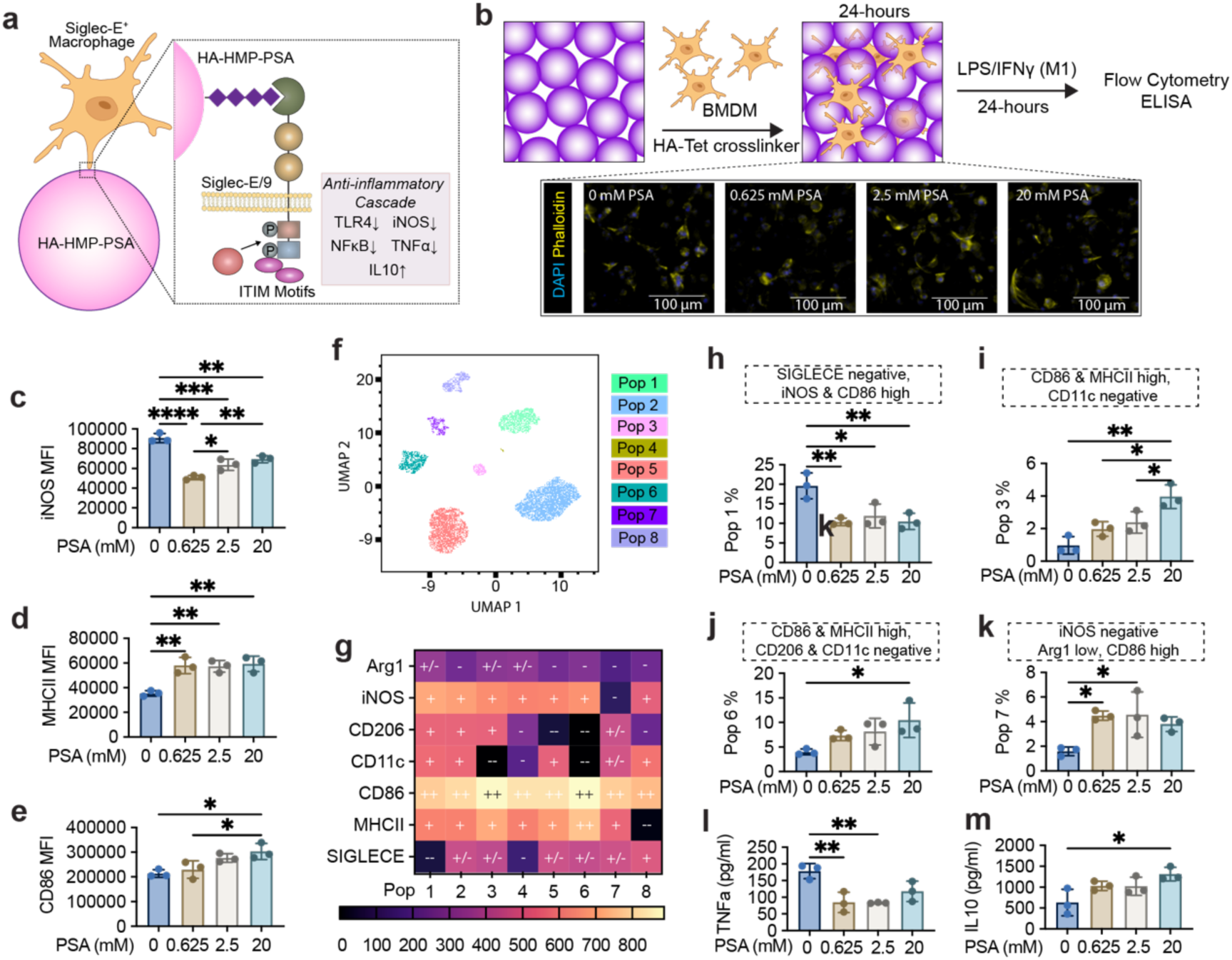
PSA-functionalized MAP scaffolds promote regulatory macrophage polarization in vitro. a) Biomimetic PSA presentation via covalent conjugation to hyaluronic acid-based hydrogel microparticles (HA-HMPs) facilitates specific interactions with Siglec-E receptors on macrophages. This interaction initiates inhibitory signaling through phosphorylation of immunoreceptor tyrosine-based inhibitory motifs (ITIMs), recruitment of phosphatases SHP- 1/SHP-2, and subsequent anti-inflammatory cascades. b) Encapsulation of BMDMs within PSA-functionalized MAP scaffolds and subsequent activation with LPS/IFNγ, with representative confocal image showing BMDMs distributed uniformly (blue: DAPI nuclei; yellow: phalloidin actin). c–e) Flow cytometry analysis of macrophage marker expression demonstrates PSA significantly decreases pro-inflammatory marker iNOS (c) and increases regulatory macrophage markers MHCII (d) and CD86 (e). f) UMAP visualization of BMDM populations reveals eight distinct macrophage clusters. g) Heatmap depicting marker expression profiles for each macrophage population cluster. h–k) Selected macrophage subsets modulated by PSA scaffolds. l–m) Cytokine analysis via ELISA confirms decreased TNFα (l) and increased IL-10 (m) secretion in PSA-treated conditions. Data represent mean ± SEM (n = 3 biological replicates; each replicate represents cells isolated from a distinct mouse). One-way ANOVA with Tukey’s post-hoc test: *p < 0.05, **p < 0.01, ***p < 0.001, ****p < 0.001.

Flow cytometry analysis revealed substantial modulation of key macrophage phenotypic markers by tethered PSA. iNOS was markedly reduced in all PSA conditions compared to controls (Figure 6c & S5). Interestingly, the greatest reduction of iNOS occurred at the lowest PSA concentration (0.625 mM), with diminishing effects at higher concentrations. This non-linear effect might reflect receptor clustering or internalization, known to occur at higher ligand densities, potentially modulating the downstream signaling efficacy and altering the expected linear dose-response relationship^[4c, 40]^. Conversely, CD86 showed a clear dose-dependent increase (Figure 6d-e). We observed no significant changes in other polarization markers (Arg1, CD206, CD11c, Siglec-E; Figure S6 a-d), indicating PSA selectively targets pathways rather than broadly modulating all M1/M2 markers. Additional analysis revealed a shift toward CD86^+^iNOS^-^ and MHCII^+^iNOS^-^ macrophage populations in PSA scaffolds, concomitant with a decrease in inflammatory iNOS^+^ subsets (Figure S7-S8). This confirms PSA’s capability to selectively suppress inflammatory polarization pathways, directing macrophages toward a more regulatory, antigen-presenting phenotype.

High-dimensional clustering revealed eight distinct macrophage subpopulations (Figure 6f). These populations varied in their marker expression patterns, highlighting phenotypic complexity beyond traditional M1/M2 classification (Figure 6g). PSA scaffolds showed a reduced proportion of macrophages with low Siglec-E alongside elevated inflammatory markers such as iNOS and CD86 (Population 1, Figure 6h). This suggests PSA’s ability to suppress particularly pro-inflammatory subsets through SIGLEC-mediated signaling^[41]^. Conversely, PSA significantly increased subsets characterized by high CD86 and MHCII expression without co-expression of CD11c or CD206 (Populations 3 and 6, Figures 6i–j), consistent with regulatory macrophage phenotypes known for enhanced antigen presentation but reduced inflammatory mediator release^[42]^. Additionally, PSA increased a distinctive population (Population 7, Figure 6k) marked by high CD86, low Arg1, and notably absent iNOS expression. The enrichment of this iNOS-negative regulatory subset aligns with the broader finding that PSA-functionalized scaffolds preferentially reduce inflammatory mediators^[16a]^.

PSA-induced polarization shifts were confirmed by cytokine analysis. PSA significantly reduced pro-inflammatory TNFα in all PSA conditions, with maximum reduction at intermediate PSA doses (Figure 6l). Anti-inflammatory IL-10 markedly increased, particularly at the highest PSA concentration (Figure 6m). This balance of cytokine modulation further demonstrates a regulatory effect exerted by PSA-MAP. Collectively, these findings illustrate that PSA-MAP scaffolds effectively guide macrophage polarization toward regulatory, anti-inflammatory phenotypes, characterized by reduced inflammatory mediator production (iNOS, TNFα) and enhanced regulatory and antigen-presenting markers (CD86, MHCII, IL-10)^[40b, 42]^.

### 2.7. PSA-MAP Modulates Immune Cell Infiltration and Polarization 48 Hours Post-Stroke *In Vivo*

We next evaluated the immunomodulatory effects of PSA-MAP scaffolds in vivo using a photothrombotic stroke model. Given robust in vitro immunomodulation observed at 20 mM PSA, this formulation was chosen for in vivo assessment (denoted as PSA_b_).

Photothrombotic stroke was induced on day 0, followed by scaffold injection into the infarct site on day 5 post-stroke (Figure 7a). Four scaffold conditions were compared: MAP-only (M), MAP with fully tethered PSA (PSA_b_), MAP with partially tethered and soluble PSA (PSA_b/s_), and MAP with fully soluble PSA (PSA_s_) (Figure 7c).

**Figure 7.**
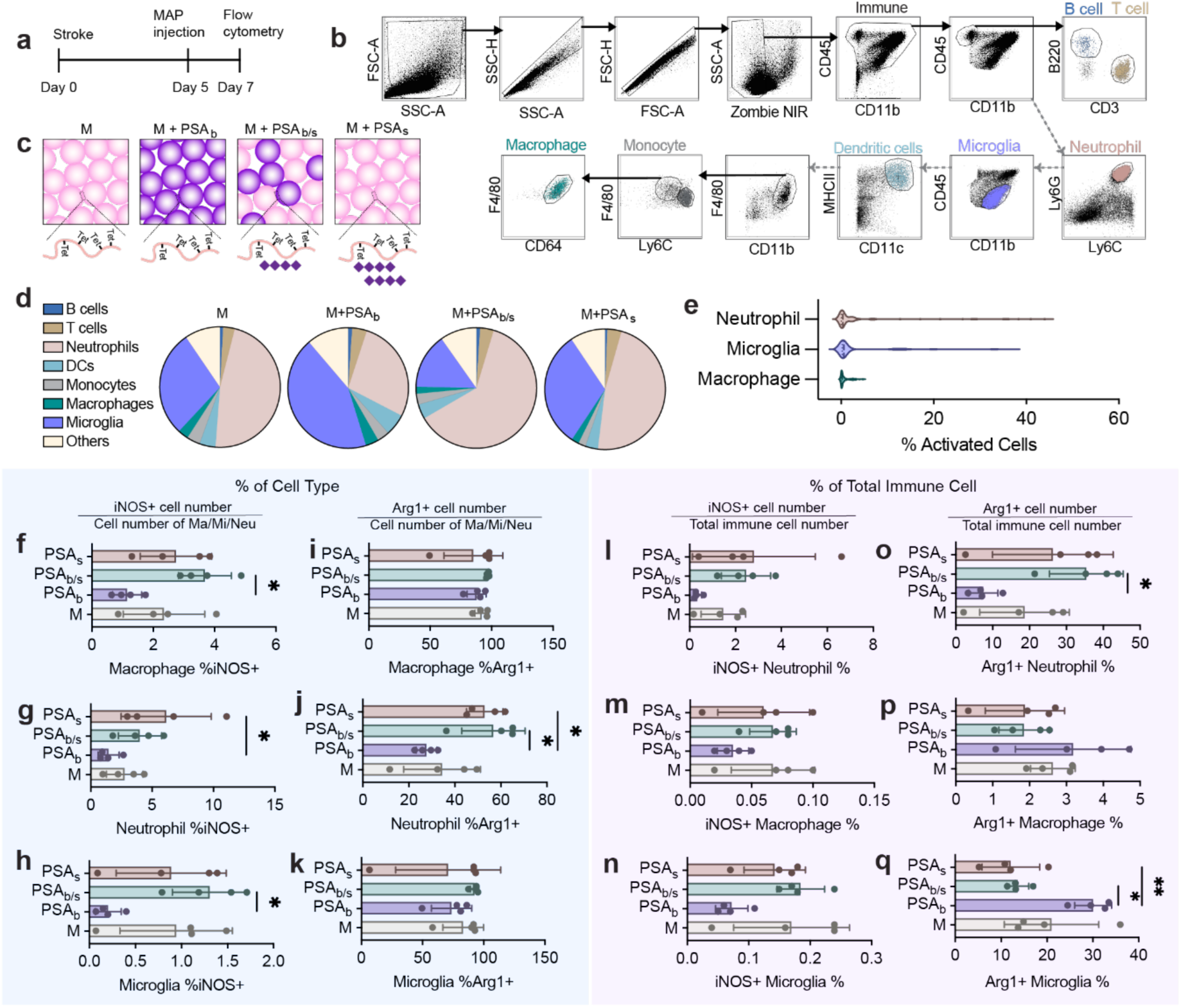
PSA-functionalized MAP scaffolds modulate innate immune profiles post-stroke in vivo. a) Experimental timeline for photothrombotic stroke induction, scaffold injection (day 5), and immune profiling by flow cytometry (day 7). b) Flow cytometry gating strategy identifying neutrophils, microglia, macrophages, dendritic cells, monocytes, T cells, and B cells. c) Schematic of four treatment conditions: MAP-only (M), MAP with fully bound PSA (PSA_b_), MAP with partially bound and soluble PSA (PSA_b/s_), and MAP with fully soluble PSA (PSA_s_). d) Pie charts illustrating immune cell composition in the infarct across treatment groups. e) Percentage of activated neutrophils, microglia, and macrophages relative to total immune cells, highlighting predominance of activated neutrophils and microglia populations. f-h) Percentage of iNOS-positive cells within (f) macrophages, (g) neutrophils, and (h) microglia, demonstrating significantly reduced pro-inflammatory iNOS expression with fully bound PSA (PSAb). i-k) Percentage of Arg1-positive cells within (i) macrophages, (j) neutrophils, and (k) microglia, showing distinct modulation of Arg1 expression across cell types by PSA treatment. l-n) iNOS-positive cell percentages among total immune cells for (l) neutrophils, (m) macrophages, and (n) microglia, confirming reduced pro-inflammatory cell abundance with PSAb. o-q) Arg1-positive cell percentages among total immune cells for (o) neutrophils, (p) macrophages, and (q) microglia, indicating significantly elevated reparative Arg1-positive macrophages and microglia with PSAb. Data presented as mean ± SEM (n=4 animal per group). One-way ANOVA with Tukey’s post hoc test: *p < 0.05, **p < 0.01, ***p < 0.001.

Flow cytometry was performed 48 hours post-scaffold injection (day 7 post-stroke) to quantify immune cell infiltration and polarization (Figure 7a–c). Infiltration by neutrophils was lower in animals treated with PSA_b_ scaffolds compared to other formulations, particularly conditions with soluble PSA (PSA_s_ and PSA_b/s_)^[43]^. Neutrophils are potent sources of reactive oxygen species (ROS), proteases, and pro-inflammatory cytokines that exacerbate neuronal injury and impede neural recovery^[44]^. Hence, the observed reduction in neutrophil recruitment suggests a beneficial anti-inflammatory shift induced specifically by the tethered PSA configuration. PSA_b_ scaffolds also increased infiltration of microglia, macrophages, and adaptive immune cells, particularly B cells (Figure 7d, S9). Among these, microglia and macrophages were specifically selected for further analysis due to their pivotal roles in orchestrating both inflammatory and regenerative processes in the post-ischemia brain^[45]^. We analyzed expression of established markers (Arg1, CD206, CD86, and iNOS) indicative of immune polarization toward pro-inflammatory or reparative phenotypes in neutrophils, microglia, and macrophages. We found that neutrophils exhibited the highest activation rates relative to total immune cell populations (Figure 7e), likely due to their abundance after stroke^[44^, ^46]^. Macrophages and microglia displayed similarly high activation frequencies within their respective populations (Figure S10), consistent with their significant involvement in the post-stroke immune response.

Phenotypic analysis revealed clear PSA-dependent modulation of inflammatory marker expression across all three cell types (Figure 7f–k). Importantly, PSA tethering markedly reduced iNOS+ neutrophil, macrophage, and microglial populations compared to soluble PSA or HA-MAP only conditions (Figure 7f–h). This suppression of iNOS expression is particularly beneficial, given that iNOS activity is strongly associated with sustained inflammation, oxidative damage, and impaired recovery post-stroke^[47]^. Conversely, soluble PSA formulations appeared to increase iNOS expression, suggesting that PSA’s immune-modulatory capacity critically relies upon tethering to the scaffold.

Intriguingly, Arg1 expression exhibited cell-type-specific modulation. In neutrophils, PSA tethering significantly reduced Arg1 positivity (Figure 7j). Although Arg1 is conventionally viewed as a reparative marker in macrophages and microglia, neutrophil-derived Arg1 has been linked to systemic immunosuppressive environments that can impair adaptive immune responses ^[48]^. Thus, reduced neutrophil Arg1 expression with PSA_b_ may reflect a favorable shift in immune balance toward more effective adaptive immunity. In contrast, macrophage and microglial Arg1 expression remained unchanged among scaffold conditions (Figure 7i, k).

High-dimensional clustering further resolved these phenotypic patterns (Supplemental Figures S11–S13). PSA tethering preferentially diminished microglial subsets expressing high CD86 and MHCII (Figure S11). Conversely, tethering of PSA subtly enriched MHCII and CD86 in macrophages, consistent with *in vitro* findings of regulatory phenotypes (Figure S12). Meanwhile, PSA tethering significantly reduced neutrophil subsets co-expressing elevated iNOS and Arg1 (Figure S13). These findings underscored the scaffold’s capacity to mitigate neutrophil activation and microglial inflammation.

Given that absolute numbers and activation percentages differ substantially among immune cell populations, we further assessed these markers as a proportion of total immune cells (Figure 7l–q). Consistent with previous analyses, PSA tethering robustly decreased overall numbers of iNOS-positive neutrophils, macrophages, and microglia (Figure 7l–n).

Importantly, PSA_b_ scaffolds significantly increased the proportion of Arg+ macrophages and microglia (Figure 7 p–q) while reducing the proportion of Arg1+ neutrophils (Figure 7o), suggesting that PSA_b_ scaffolds minimize pro-inflammatory neutrophil activation while promoting reparative macrophage and microglial phenotypes^[43^, ^49]^. CD86 and CD206 displayed minor variations among conditions (Figure S14), emphasizing the unique sensitivity of iNOS and Arg1 as functional biomarkers of immune polarization in this therapeutic context.

Although differences at early time points primarily distinguished PSA_b_ from soluble PSA scaffolds, fewer differences were observed between PSA_b_ and MAP-only scaffolds, likely due to complex inflammatory dynamics post-stroke. Nonetheless, PSA_b_ scaffolds consistently trended toward lower pro-inflammatory iNOS marker expression and enhanced regulatory immune populations. Interestingly, in vivo soluble PSA formulations exhibited unintended pro-inflammatory responses, likely due to uncontrolled diffusion and broader receptor interactions. In contrast, tethered PSA enabled localized receptor engagement, suggesting PSA presentation is critical for achieving effective immunomodulation *in vivo*.

Together, these results demonstrate that PSA_b_ scaffolds modulate immune cell infiltration and polarization *in vivo*, characterized by reduced neutrophil-driven inflammation, decreased pro-inflammatory marker expression (iNOS, TNFα), and increased reparative Arg1 immune populations. Given the temporally dynamic inflammatory environment post-stroke, we subsequently investigated immunomodulatory effects at day 15 post-stroke.

### 2.8. PSA-MAP Limits Peri-infarct Inflammation and Enhance Resident Microglia at Subacute Stages

Having established that PSA-MAP scaffolds modulate innate immune responses shortly after injection, we next examined immune cell dynamics at day 15 post-stroke (10 days post-scaffold injection), utilizing immunofluorescence (IF) staining to characterize spatial patterns of immune cells within infarct and peri-infarct regions (Figure 8a). Given the pivotal roles of macrophages and microglia in sustained inflammation and reparative processes, understanding their localization within infarct and peri-infarct regions at this later stage is crucial for evaluating sustained therapeutic outcomes and tissue repair potential. Given our earlier data demonstrating the superiority of conjugated PSA (PSA_b_) scaffolds in modulating beneficial immune responses, we specifically compared the PSA_b_ condition with the MAP-only (HA) control at this later timepoint.

**Figure 8.**
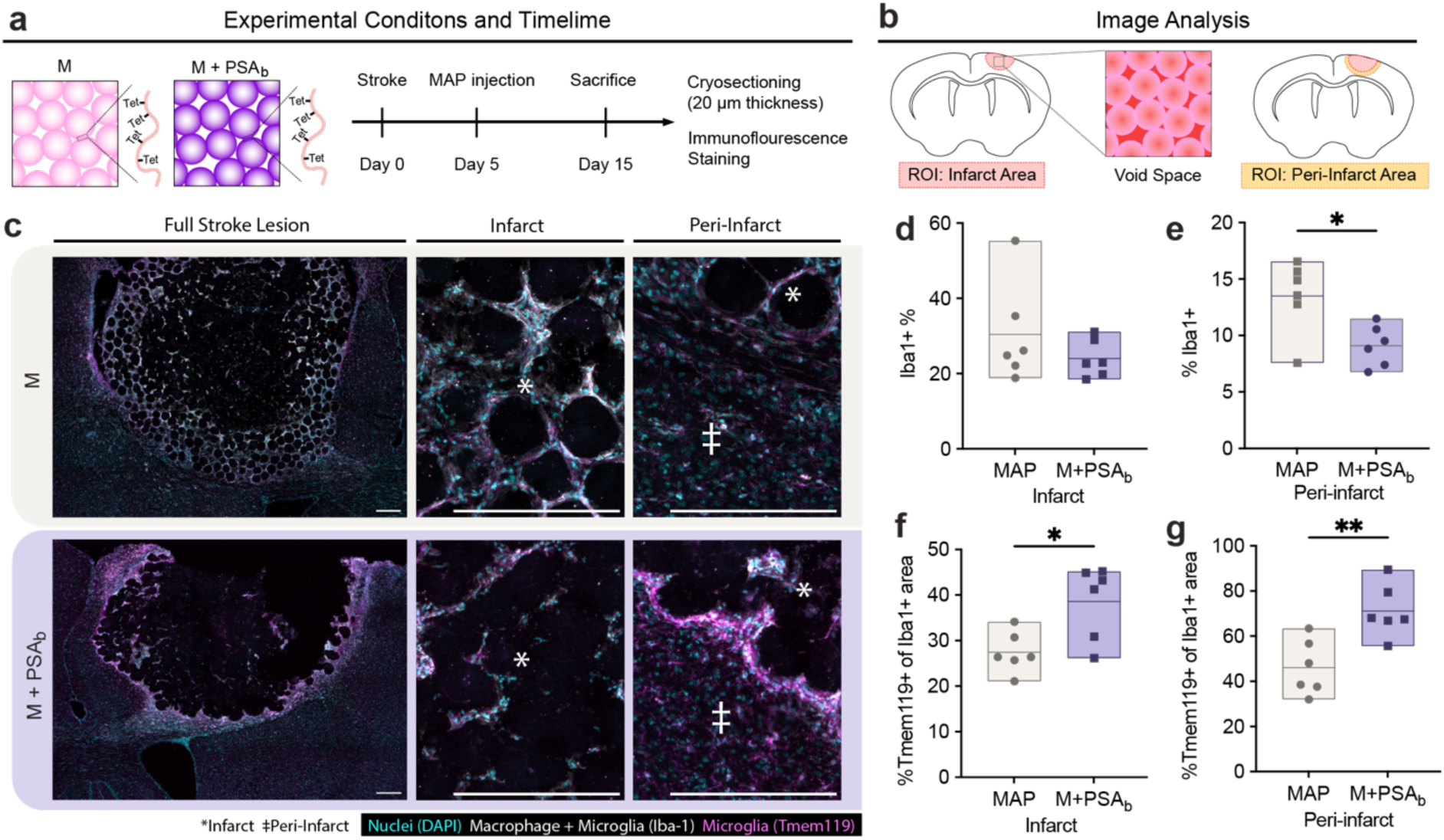
PSA-functionalized MAP scaffolds support beneficial microglial response at late subacute stage post-stroke. a) Timeline depicting photothrombotic stroke induction, scaffold injection (day 5), and brain tissue immunofluorescence analysis at day 15. b) Representative immunofluorescence images (Iba1, TMEM119, and DAPI) comparing infarct and peri-infarct regions between MAP-only (m) and fully bound PSA-MAP (PSA_b_) scaffolds. d-d) Quantification of Iba1-positive area (% of total nuclei area) within (c) infarct and (d) peri- infarct regions, highlighting decreased microglia/macrophage presence in peri-infarct areas with PSA_b_. e-f) Percentage of TMEM119-positive area within the Iba1-positive regions in (e) infarct and (f) peri-infarct areas, showing significantly increased proportion of resident microglia with PSA_b_ scaffolds. Each data point is a biological replicate averaged from two coronal sections and plotted in a floating bar (n = 6). Two-tailed, unpaired Student’s t-test was performed: *p < 0.05, **p < 0.01. Scale bars represent 200 µm.

IF staining revealed distinct abundance of microglia and macrophages (Iba1+ cells) within infarct and peri-infarct areas (Figure 8 b-c). In the infarct, there were no significant differences between the PSA_b_ and MAP groups, with both conditions showing approximately 23% of lesion area coverage by Iba1+ cells (Figure 8d). This is consistent with ongoing clearance and repair processes required during the subacute phase of stroke recovery^[43]^. However, pronounced differences emerged in the peri-infarct: PSA_b_ scaffolds significantly reduced total Iba1+ cell coverage compared to controls (Figure 8e). This likely reflects a reduction in persistent inflammation and may lead to an environment that is conducive to neural repair^[43]^.

To further distinguish between microglia and macrophages within the Iba1+ population, we next examined TMEM119, a selective resident microglia marker. PSA_b_ scaffolds demonstrated significantly higher proportions of TMEM119+ microglia relative to the total Iba1+ population in both infarct and peri-infarct regions (Figure 8f-g). This finding suggests enhanced retention of resident microglia and reduced peripheral macrophage infiltration. Given that resident microglia typically adopt phenotypes associated with tissue repair and homeostasis rather than inflammatory activation, the observed enrichment of resident microglial populations represents a favorable immunomodulatory outcome^[45a, 50]^.

These findings extend our earlier observations at day 7 post-stroke and confirm that PSA_b_ scaffolds sustain beneficial microglial responses into later stages of stroke recovery. By preferentially supporting resident microglial populations and reducing the recruitment of peripheral macrophages, PSA-MAP scaffolds foster a microenvironment conducive to endogenous repair processes ^[43]^.

## 3. Conclusion

In this study, we introduced a biomimetic platform utilizing PSA-functionalized MAP scaffolds to modulate immune responses following ischemic stroke. The PSA-MAP platform mimics the physiologic presentation of PSA by covalently tethering it via the reducing end.

We demonstrated successful endotoxin removal from PSA, maintenance of scaffold biocompatibility after PSA incorporation, and confirmed PSA’s potent immunoregulatory activity among various SA derivatives tested. Chemical characterization established a robust functionalization process, preserving desired scaffold mechanics, while oxidation of the PSA non-reducing end further enhanced enzymatic stability. By encapsulating BMDMs within PSA-MAP scaffolds, we identified significant shifts towards regulatory immune phenotypes (reduced iNOS/TNF-α; increased CD86/MHCII/IL-10). Clustering analysis reinforced these observations, highlighting a decrease in pro-inflammatory macrophage subsets and enrichment of regulatory macrophage populations^[40b, 51]^.

Injection with PSA-MAP in a murine ischemic stroke model dramatically modulated the immune landscape. Neutrophils were significantly diminished in number and activation status in the presence of tethered PSA^[44]^. Conversely, while total Arg1+ neutrophils were also reduced in PSA scaffold conditions, this decrease is beneficial given that neutrophilic Arg1 expression often correlates with disrupted T-cell immunity and ineffective tissue repair in peripheral system^[48b, 52]^. Consequently, the observed PSA-mediated suppression of neutrophil infiltration and activation may be a substantial therapeutic advantage, promoting an environment conducive to reparative processes.

In parallel, PSA-MAP scaffolds enhanced reparative phenotypes within resident microglia and infiltrating macrophages, characterized by a substantial increase in Arg1 expression. This coordinated modulation—reducing detrimental neutrophil iNOS activity while simultaneously enhancing Arg1+ microglia/macrophages—reflects a targeted immunological balance critical for effective resolution of inflammation and initiation of repair pathways^[49a]^. In addition, beneficial immune alterations persisted into a later subacute stage, indicated by decreased overall inflammatory cell burden, particularly within peri-infarct zones, accompanied by enrichment of resident microglia populations. Collectively, our study establishes PSA-functionalized MAP scaffolds as a promising approach for targeted immunomodulation during critical periods of post-stroke tissue recovery. By effectively dampening harmful neutrophil-driven inflammation and promoting reparative macrophage/microglia responses through strategic modulation of Arg1 and iNOS pathways, this biomaterial system offers significant translational promise for enhancing intrinsic brain repair processes and improving outcomes following ischemic stroke.

## 4. Experimental Section/Methods Primary Murine Macrophages Culture

Bone marrow-derived macrophages (BMDM) were isolated from 8–12 weeks old C57BL/6 (female) in accordance with institutional and state guidelines and approved by the Duke University’s Division of Laboratory Animal Resources (DLAR). Animals were anesthetized, and the tibias and femurs were collected. The bone marrow was flushed out from the bones and broken into cell suspension by repeated pipetting. The cells were differentiated in a culture medium (10% v/v heat-inactivated fetal bovine serum and 1% v/v antibiotic-antimycotic in Iscove’s Modified Dulbecco’s Medium, Gibco) with 15 ng mL^−1^ macrophage colony-stimulating factor (Peprotech). The medium was changed on days 1, 4, and 8. After full maturation, at day 10 the cultured cells were tested for murine macrophage pan marker CD11b and F4/80 by flow cytometry to confirm the macrophage percentage (CD11b+F4/80+).^[53]^

### Macrophage In Vitro Culture on Tissue Culture Plastics

BMDM were detached from the culture flask on day 10 with 0.1 X TrypLE + 1mM EDTA + 1% antibiotic-antimycotic made from 10 X TrypLE Select solution (Thermo Fisher). The cells were then plated at 52k cells cm^-2^ onto 12-well non-treated plates (VWR) and cultured in 15 ng mL^−1^ M-CSF media for overnight before 24-hr cytokine activation. M1 activation was with 20 ng mL^−1^ LPS (Thermo Fisher) and IFN-*γ* (Peprotech), and M2 activation was with 20 ng mL^−1^ IL-4 (Peprotech).^[54]^ For conditions involving sialic acid derivatives, each derivative was added concurrently with activation cytokines. To standardize dosing, each sialic acid derivative—regardless of its structural complexity (monomeric, polymeric, or glycoconjugate form)—was weighed based on the molecular weight of the N-acetylneuraminic acid (Neu5Ac) monomer. Specifically, for polysialic acid (PSA), composed entirely of Neu5Ac units linked by α2,8-glycosidic bonds, the total amount of Neu5Ac monomer was directly calculated from the polymer mass. Similarly, for CMP-Neu5Ac, 3’- sialyllactose (3’SL), and 6’-sialyllactose (6’SL), which possess defined chemical structures, the Neu5Ac content was directly determined based on the known molecular weights. All sialic acid derivatives were thus normalized to a uniform final concentration of 5 × 10⁻³ M Neu5Ac monomer units.

### Real Time RT-qPCR of BMDM

Total RNA from BMDM cultured on tissue culture plastics was isolated by RNeasy Mini Kit (QIAGEN) following the manufacturer protocol. The concentration and purity of the total RNA were determined using a Nanodrop spectrophotometer (Thermo Fisher Scientific).

Total RNA (1μg) was transcribed into first strand cDNA using an iScript cDNA synthesis kit (Bio-Rad) according to the manufacturer’s procedure. Expression of SIGLEC-E and GAPDH mRNAs were measured by quantitative PCR (qPCR) using the appropriate primers with an iTaq Universal SYBR Green Supermix (Bio-Rad) on a StepOnePlus Real-Time PCR System (Applied Biosystems). The first cycle was 30 seconds at 95 °C, and the 40 subsequent cycles were at 95 °C for 15 s and 60 °C for 1 min. Relative expression levels were calculated using the ΔΔCT method (2^−ΔΔCT^) using GAPDH was used as a reference gene. The primers used in the experiment are: GAPDH, forward 5’- TGTCCGTCGTGGATCTGAC-3’ and reverse 5’- CCTGCTTCACCACCTTCTTG-3’; Siglec-E (primer 1), forward 5’- GTCTCCACAGAGCAGTGCAACTTTATC-3’ and reverse 5’- TGGGATTCAACCAGGGGATTCTGAG-3’; Siglec-E (primer 2), 5’- TGGTACAGGGAAGGAACCGA-3’ and reverse 5’-GTGAGGGCTGTTACAACCAGA-3’.

### Endotoxin Test

Endotoxin levels were quantified with the Pierce Limulus Amebocyte Lysate (LAL) Chromogenic Endotoxin Kit (Thermo Fisher Scientific) according to the manufacturer’s instructions. High-molecular-weight hyaluronic acid (HA) was obtained from Contipro, and polysialic acid (PSA) was sourced from two commercial suppliers: Santa Cruz Biotechnology (vendor 1, MW range approximately 24 000 – 38 000 Da as specified by the vendor) and BOC Sciences (vendor 2, MW range approximately 50 000 – 60 000 Da as specified by the vendor). Although these PSA samples differ in the molecular weight ranges provided by the respective vendors, they are chemically identical, composed of Neu5Ac repeat units connected by α2,8 glycosidic linkages.

### Modification of Hyaluronic Acid with Norbornene

Hyaluronic acid–norbornene (HA–NB) was synthesized through the activation and subsequent functionalization of the HA carboxylic acid group by dissolving 1.0 g of HA (MW 79 000 Da reported from the vendor) (Contipro) and 3.1 g 4-(4,6- dimethoxy[1.3.5]triazin-2-yl)-4-methylmorpholinium chloride (DMTMM) (MW: 294.74 Da) (TCI America, Portland, OR, USA) (4 molar equivalents) each in 40 mL of 200 × 10^−3^ M MES buffer, pH 5.5, combining the solutions and allowing the reaction to stir for 10 min. Then 0.677 mL of 5-norbornene-2-methanamine (a mixture of isomers) (NMA) (TCI America, Portland, OR) (2 molar equivalents) was added dropwise into the mixture. The reaction was stirred at room temperature overnight and then precipitated in 1 L of 100% Ethanol at 4 °C. All precipitates were collected and dissolved in 2 M brine solution and dialyzed against DI water for 30 min then 1 M brine solution for 30 min. This dialysis process was repeated three times and then dialyzed against DI water for 24 h. The final solution was collected and lyophilized to yield the final product. HA–NB was confirmed by ^1^H-NMR with 31% - 45% NB functionalization. ^1^H NMR resonances of pendant norbornenes in D_2_O, δ6.33 and δ6.02 (vinyl protons, endo), and δ6.26 and δ6.23 ppm (vinyl protons, exo) were compared to the resonance of the HA methyl group δ2.05 ppm to determine the extent of functionalization (Figure S15). All equivalents were based on the moles of the HA repeat unit.

### Modification of Polysialic Acid Reducing End with Thiol and Tetrazine

PSA–SH was prepared by reductive amination adapted from previously described method.^[28]^ Polysialic acid (MW 24000 – 38000) (Santa Cruz Biotechnology) after endotoxin removal was dissolved at 50 mg/mL and cystamine dihydrochloride (Chem-Impex) dissolved at 12.4 mg/mL in 0.1 M odium borate buffer with 1 M NaCl, pH 8.5, and stirred at 40 °C for 2 h to promote ring-opening of the terminal Neu5Ac monomer at the reducing end and formation of Schiff base between the amine of cystamine dihydrochloride and the ketone at C2 of the terminal Neu5Ac monomer. NaBH3CN was then added to the solution at a final concentration of 300 mM and reacted at 55 °C for 96 hours. The reaction mixture was incubated with 150 mM dithiothreitol (DTT) for 12 h to introduce a free thiol group, dialyzed (MWCO: 3.5 kDa) against DI water for 30 min then 1 M brine solution for 30 min. This dialysis process was repeated three times, dialyzed against DI water for 24 h and lyophilized for 3 days. Initially, we considered direct conjugation of PSA-thiol (PSA-SH) to HA-norbornene (HA-NB) microparticles through thiol–norbornene click chemistry. However, preliminary rheological experiments revealed that unmodified PSA could directly react with HA-NB in the presence of LAP photoinitiator and UV irradiation (*λ* = 365nm), independent of any added di-thiol crosslinker. Specifically, solutions of HA-NB mixed solely with PSA (without any crosslinker) were subjected to UV exposure (initiated at 1 minute) in the presence or absence of LAP, and the storage modulus (G’) was monitored. The addition of LAP resulted in a rapid and significant increase in G’ (1000–3000 Pa), indicating unintended crosslinking reactions occurring directly between PSA and HA-NB under UV conditions (Figure S16). Without LAP, G’ remained consistently low, confirming the photoinitiator- dependent nature of this unintended crosslinking. Due to uncertainty about the exact chemistry and resulting glycan orientation of these unintended products, we opted to pursue an alternative route involving the controlled conversion of PSA-SH to PSA-tetrazine (PSA- Tet) for subsequent bio-orthogonal conjugation to HA-NB, ensuring precise chemical control and physiological mimicry. PSA-SH-Tet was prepared by thiol-maleimide Michael addition. PSA-SH was dissolved in 4.5 x 10^−2^ mM HEPES solution, pH 7.4, at 6.66 mg/mL and treated with TCEP at 2.78 x 10^−4^ M for 15 min at room temperature to reduce disulfide bonds.

Methyltetrazine-PEG_4_-maleimide then was added to the solution at a final concentration of 1 mg/ml and reacted at room temperature for 1 h to allow the reaction between the thiol and the maleimide. The solution was dialyzed (MWCO: 3.5 kDa) against DI water for 30 min then 1 M brine solution for 30 min. This dialysis process was repeated three times, dialyzed against DI water for 24 h and lyophilized for 3 days. Successful functionalization of tetrazine was verified with NMR (Figure S17). The presence of characteristic proton signals at δ8.35 (2H) and δ7.20 (2H), corresponding to the aromatic protons of the tetrazine moiety in D_2_O, confirmed successful tetrazine conjugation.

### Oxidization of Polysialic Acid Non-Reducing End

The non-reducing end of PSA-SH or PSA-SH-Tet was activated by selective periodate oxidation as described previously.^[29^, ^55]^ PSA polymer (reducing end modified) was dissolved at 40 mg mL^-1^ and mixed with freshly prepared 2 x 10^-1^ mM sodium metaperiodate stock solution (Sigma-Aldrich) to achieve a molar ratio of sodium periodate to vicinal diol of 1.1:1 in 4.5 x 10^-2^ mM HEPES buffer. The solution was stirred 4 °C for 4 h in the dark to allow oxidation of the vicinal diol at the non-reducing end. To stop the reaction, ethylene glycol was added to the reaction mixture at a molar ratio of ethylene glycol to sodium periodate of 0.2:1 and left to stir at 20 °C for 30 min. The polymer is now modified with an aldehyde at the terminal Neu5Ac unit on its reducing end. Aniline catalyzed oxime ligation is then performed to conjugate either biotin or AF-568 to the reducing end, as described previously.^[29]^ AZDye 568 Hydroxylamine (Vector Laboratories) or Biotin-PEG3-oxyamine (BroadPharm) was dissolved in H_2_O at 5 x 10^-2^ mg/μL and added to the reaction mixture at a molar ratio of oxyamine to vicinal diol of 1.5:1 and stir for 90 min at room temperature. The final Biotin-PSA-Tet or AF568-PSA-Tet solution was dialyzed (MWCO: 3.5 kDa) against DI water for 30 min then 1 M brine solution for 30 min. This dialysis process was repeated three times, dialyzed against DI water for 24 h and lyophilized for 3 days. Successful functionalization of AF568 was verified using ^1^H NMR spectroscopy by comparing the spectra of AF568-PSA-Tet and free AZDye568-hydroxylamine (Figure S18). The presence of characteristic aromatic and alkene proton peaks at δ 5.8 ppm in both spectra confirmed successful conjugation of AF568 to the polymer backbone.

### Diffusion-Ordered NMR Spectroscopy (DOSY)

Polymer molecular weights were determined using DOSY via previously reported methods.^[56]^ Briefly, DOSY experiments were performed on a Bruker Avance III 700 MHz spectrometer equipped with a Broadband Observe (BBO) probe. Polymer samples were diluted (10 mg/mL) in deuterium oxide and experiments were performed at 25 °C. Measurements were taken using the TopSpin3.5pl5 *ledbpgp2s* pulse program with sine-shaped gradients ramped linearly over 25 scans with a first gradient amplitude of 2% and a final gradient amplitude of 95%. The gradient duration (P30) and diffusion delay (D20) were adjusted for each temperature to achieve ≥ 90% signal reduction with the final gradient. Collected spectra were phase- and baseline-corrected and analyzed using the DOSY Peak Fit Auto Decay Component model on MestReNova (v 14.2.0-26256). Polymer diffusion coefficients were calculated using the median of the values collected for ^1^H NMR resonances with clear signals at 95% gradient amplitude. Relative weight-average molecular weights (*M_w_*) of the polymers were calculated from PEG calibration curves previously developed (Figure S2).^[56a]^

The percent chain cleavage was calculated based on the reduction in the average degree of polymerization (DP_n_) measured by DOSY NMR after each conjugation step, using the following equation: % chain cleavage = (DP_initial_ – DP_final_) / DP_initial_ x 100%, where DP_initial_ and DP_final_ represent the average degrees of polymerization before and after each chemical conjugation step, respectively.

### HMP Production and Purification

HA–NB hydrogel microparticles (HMPs) were produced using a planar flow-focusing microfluidic device to create uniform particles (Figure 3A). A 1 mL gel precursor solution was made by dissolving HA–NB in 3 × 10^−1^ M HEPES pH 7.5, di-thiol MMP sensitive linker peptide (Ac-GCRDGPQGIWGQDRCG-NH2, Genscript) (SH/HA ratio of 14), tris(2- carboxyethyl)phosphine (TCEP) (Sigma-Aldrich) (TCEP/SH ratio of 0.25), 5 x 10^−4^ M RGD peptide (Ac-RGDSPGERCG-NH2, Genscript), and 9.90 × 10^−3^ M lithium phenyl(2,4,6- trimethylbenzoyl)phosphinate photoinitiator (LAP) (TCI America, Portland, OR, USA). The final HA–NB in the precursor solution should be at 3.4% (w/v). The solution was filtered through a 0.22 μm sterile filtered before being transferred to a 1 mL BD Leur-Lok syringe and connected to the inner inlet of the microfluidic device using tubing, resulting in approximately 30% solution during the filtering process. A 5 mL BD Leur-Lok syringe was filled with 5% (v/v) Span-80 in heavy mineral oil and attached to the outer inlet of the microfluidic device. A single syringe pump was used to push the differently sized syringes at asymmetric flow rates at a flow rate ratio of ≈ 6.4:1 (oil:aqueous). The precursor solution was pinched at the flow focusing region by the oil and the HMP droplets were crosslinked by consistent exposure to UV light (20 mW cm^−2^) off chip using an OmniCure LX500 LED Spot UV curing system controller with a OmnniCure LX500 LED MAX head (365 nm, 40% power). A 15 mL conical tube wrapped in foil was used to collect the microgel emulsion.

Upon completion of fabrication, the suspension was centrifuged at 5250 rcf for 5 min and the supernatant oil was aspirated off in a sterile hood. Everything following was performed under sterile conditions. The HMPs were then washed with sterile filtered HEPES buffer and centrifuged again at 5250 rcf for 10 min. Washing was repeated several times until no more oil was visible in the supernatant. Following washes, endotoxin tests were performed. HMP endotoxin levels were consistently below 0.2 endotoxin U mL^−1^. HMPs were then stored at 4 °C until further use.

### HMP Post-Fabrication Modification with PSA-SH-Tet

HA–NB HMPs were modified post-fabrication via inverse electron demand Diels–Alder tetrazine–norbornene click reaction, in which excess norbornene groups on the HMPs were functionalized with AF555–PSA-Tet to fluorescently tag the HMP and determine post-modification efficiency. For PSA conjugation efficiency quantification, the HMPs were incubated in 3 × 10^−1^ M HEPES pH 7.4 containing AF555–PSA-Tet at initial Neu5Ac monomer concentrations of 0 M, 6.25 x 10^-4^ M, 1.25 x 10^-3^ M, 2.5 x 10^-3^ M, 5 x 10^-3^ M, 1 x 10^-2^ M, or 2 x 10^-2^ M at 37 °C for 4 h with agitation every 1 h, as previously noted. The suspended HMPs were then pelleted by centrifuging at 14 000 rcf for 15 min. The HMPs were washed three times with 3 × 10^−1^ M HEPES and recovered with the same centrifugation conditions. Modified HMPs were imaged on a Nikon Ti Eclipse with a 20× air objective. For *in vitro* cell culture and *in vivo* injection, HA–NB HMPs were modified incubated in 0.22µm sterile filtered 3 × 10^−1^ M HEPES pH 7.4 containing PSA-SH-Tet at final Neu5Ac monomer concentrations of 0 M, 1.25 x 10^-3^ M, 2.5 x 10^-3^ M, or 2 x 10^-2^ M at 37 °C for 4 h with agitation every 1 h. The suspended HMPs were then pelleted by centrifuging at 14 000 rcf for 15 min. The HMPs were washed three times in a laminar flow tissue-culture hood with sterile filtered 3 × 10^−1^ M HEPES, one time wash with 70% EtOH for 8 min, three times again with sterile filtered 3 × 10^−1^ M HEPES, and recovered with the same centrifugation conditions as above. The recovered HMPs were then exposed to UV sterilization for 2 hours before in *vivo* and *in vitro* applications.

### Cellpose Quantification of Fluorescent PSA Conjugation Efficiency

Fluorescence-PSA-conjugated hydrogel microparticles (HMPs) confocal images were pre- processed using Gaussian blur (kernel size 5×5) and bilateral smoothing (diameter 9, sigmaColor 75, sigmaSpace 75) to enhance segmentation quality. Segmentation was then performed using the Cellpose Cyto3 model version 3.0.10 with a flow threshold of 0.8, a cell probability threshold of −1.0, and a fixed object diameter of 91 pixels.^[57]^ Non-gel artifacts were excluded based on object size and FITC signal intensity, corresponding to hyaluronic acid (HA) autofluorescence. Conjugation efficiency of PSA–tetrazine was quantified by calculating the mean AF568 fluorescence intensity within each segmented hydrogel. These intensity values were subsequently used to generate a standard curve for PSA conjugation.

### Modification of Hyaluronic Acid with Tetrazine

HA-tetrazine (HA-Tet) synthesis was adapted from previously reported methods.^[58]^ using 79 kDa HA with molar equivalents of 1:1:0.07 of HA-repeat units to 4-(4,6-dimethoxy-1,3,5-triazin-2-yl)-4-methylmorpholinium chloride (DMTMM, Thermo Fisher Scientific) to tetrazine-amine (Chem-Impex) to successfully modify approximately 4-6% of HA repeat units with tetrazine. All polymer modifications were purified with dialysis (12–14 kDa molecular weight cut-off) and subsequently quantified by ^1^H NMR. ^1^H NMR resonance shifts of pendant tetrazine groups at δ8.5 (2H) and δ7.7 (2H) (aromatic protons) were compared to the resonance of the HA methyl group δ2.05 ppm to determine functionalization (Figure S19).

### Generation of Scaffolds and Rheological Characterization

MAP scaffolds were assembled via inverse-electron-demand Diels–Alder tetrazine– norbornene click reaction, in which excess norbornene groups on microgels were annealed by HA-Tet to form tetrazine-mediated MAP (Tet-MAP) scaffolds. The elastic property of the annealed Tet-MAP scaffolds was measured as the storage modulus (*G*′) and loss modulus (*G*′′) using a parallel plate rheometer (Anton Paar, Physica MCR 301). The Tet-MAP scaffolds were formed by combining and mixing packed HMPs with HA-Tet at the desired HA-Tet/HA–NB ratio of 0.35. The solution (40 μL) was then placed between two glass slides coated with Sigmacote (Sigma-Aldrich), a solution of chlorinated organopolysiloxane in heptane that forms a hydrophobic film on glass. The slides were separated by 1 mm spacers on either side, secured with binder clips, and incubated for 1 h at 37 °C unless stated otherwise. This method was used to create 8 mm by 8 mm height discs of the Tet-MAP scaffolds for rheological testing. The crosslinked scaffold was transferred into PBS overnight to allow for swelling. Amplitude sweep was first performed on the crosslinked scaffolds using an angular frequency of 2.5 rad s^−1^ with a sheer strain range of 0.1% to 100% to determine the linear viscoelasticity range. Frequency sweep was performed on the scaffolds using a sheer strain of 1% with an angular frequency range of 0.1 to 100 rad s^−1^. The storage and loss moduli were reported from averaging the values between 1 rad s^−1^ and 10 rad s^−1^ from these frequency sweep measurements. To evaluate the stress relaxation behavior of the Tet-MAP scaffolds, the samples were placed under 10% strain and the stress was measured over a period of 24 hours.

### Neuraminidase Degradation of PSA

MAP scaffolds were seeded into a custom polydimethylsiloxane (PDMS) MAP-seeding chamber fabricated as previously described and then incubated with bacterial neuraminidase to assess polysialic-acid degradation^[59]^. Briefly, the MSC’s negative mold was produced on a Form 2 stereolithography 3-D printer (Formlabs, Inc.; Duke SMIF). Each well features a 4 mm-diameter × 1 mm-high culture cavity capped by a 5 mm-diameter × 3 mm-high media reservoir that holds ≥ 50 µL. The mold was cast in PDMS, cured, and the finished wells were plasma-bonded to glass coverslips. HA hydrogel microparticles (HA-HMPs) were first functionalized with either PSA-SH-Tet (non-oxidized, “PSA-HMP”) or biotin-PSA-SH-Tet (oxidized, “Biotin-PSA-HMP”) at an initial Neu5Ac monomer concentration of 2 x 10^-2^ M, following the protocol detailed above. After conjugation, HMPs were washed, pelleted at 14 000 × g for 15 min, the supernatant discarded, and the spin–decant cycle repeated once more to achieve a tightly packed pellet. The packed HMPs were then mixed with HA-Tet solution (HA-Tet/HA-NB ratio = 0.35), mixed thoroughly on ice, and 20 µL of the suspension was dispensed into each MAP-seeding chamber to create a hemispherical dome. Constructs were annealed at 37 °C for 1 h and then rehydrated with 1X PBS before use. For PSA-MAP degradation assays, α2-3,6,8 neuraminidase (New England Biolabs) was diluted in DMEM/F- 12 (Gibco) supplemented with 1 % (v/v) penicillin–streptomycin, 5 % (v/v) fetal bovine serum (R&D Systems) and 1X Glyco Buffer to final activities of 4 U µL⁻¹ (high) or 0.2 U µL⁻¹ (low). After gently aspirating PBS from each MAP-seeding chamber using a 100 µL pipette, 100 µL of the enzyme solution was added, and the scaffolds were incubated at 37 °C for either 4 h or overnight (∼16 h). Plates containing the PDMS wells were paraffin-sealed and housed with a PBS reservoir to prevent evaporation. At each endpoint (4 h & 16 h), the supernatant solution was collected and stored at –20 °C for subsequent DMB labeling. To assess enzymatic degradation of soluble polysialic acid, reaction mixtures (50 µL) were prepared containing either non-oxidized PSA-SH-Tet or oxidized biotin-PSA-SH-Tet at a Neu5Ac monomer concentration of 500 µM. α2-3,6,8 neuraminidase (New England Biolabs) was added to each mixture to a final activity of 4 U µL⁻¹ in H₂O supplemented with 1× GlycoBuffer. Reactions were incubated at 37 °C overnight (∼16 h), heat-inactivated at 65 °C for 10 min, and stored at –20 °C until DMB labeling.

### DMB Labeling and HPLC Quantification of Neu5Ac

Neu5Ac released from either soluble polysialic acid solutions or MAP-scaffold supernatants was quantified with the Agilent AdvanceBio Sialic Acid Profiling & Quantitation Kit (formerly ProZyme) by derivatizing sialic acids with 1,2-diamino-4,5- methylenedioxybenzene (DMB) according to the manufacturer’s protocol. Matrix-matched calibration curves were prepared for each sample type: soluble-PSA standards of 100, 200, 300, 400, and 500 µM Neu5Ac were made in water containing 1 × GlycoBuffer, while MAP-supernatant standards of 5, 10, 50, 100, and 200 µM Neu5Ac were prepared in DMEM/F-12 supplemented with 5 % (v/v) fetal bovine serum, 1 % (v/v) penicillin-streptomycin, and 1 × GlycoBuffer. After DMB labeling, samples were analyzed on an Agilent 6224 LC-MS/TOF equipped with a Poroshell 120 EC-C18 column (3 × 50 mm, 2.7 µm) held at 40 °C, using water and 5-15% gradient acetonitrile as mobile phases, UV detection at 370 nm, positive-ion electrospray MS detection, and an injection volume of 5–10 µL. Soluble-PSA standards and samples were diluted ten-fold with water before injection, whereas MAP-supernatant samples and standards were injected neat. Neu5Ac concentrations were determined by integrating the DMB-Neu5Ac peak and referencing the appropriate standard curve (Figure S20).

### Computational Analysis of MAP Scaffolds

HA-HMPs and PSA-HMPs were modified post-fabrication with Sulfo-Cyanine3 tetrazine (Lumiprobe) at a 0.0133 mM final concentration as described above to fluorescently tag the HMPs. HA-HMPs and PSA-HMPs are mixed with HA-Tet solution (HA-Tet/HA-NB ratio = 0.35), mixed thoroughly on ice with 3 compositions: 100% HA-HMPs, 50% HA-HMPs with 50% PSA-HMPs, and 100% PSA HMPs, and 20 µL of the suspension was dispensed into a MAP-seeding chamber to create a hemispherical dome, incubated at 37 <u>°</u>C for 1 h and rehydrated with 1X PBS for 1 hr. Fluorescence imaging of the scaffolds was performed using a Nikon Ti Eclipse microscope equipped with a C2 laser with a 20× air objective at a 2 µm step-size. Confocal z-stack images of each scaffold were first converted into a data format that lists the 3D-voxels associated with each unique particle using a custom algorithm.

Processed data was then fed into LOVAMAP, and 3D pores were identified, which represent open 3D spaces between particles^[35]^. Data were reported for all 3D pores – both interior and those that lie at the surface of the sample. To characterize the void space of the materials, distributions were reported for 3D pore volume (pL), pore surface area (µm^2^ 1000^-1^), pore characteristic length (µm), aspect ratio, pore average internal diameter (µm), pore longest length (µm) (Figure S4).

### Macrophage In Vitro Encapsulation in MAP Scaffolds

BMDM were detached from the culture flask with TrypLE express enzyme solution (Thermo Fisher). The cell pellet was prepared at a final concentration of 10 000 cells µL^-1^ of MAP scaffolds. Subsequently, the HMP mixture (50 µL) containing HA-Tet (HA-Tet/HA-NB ratio = 0.35) was added to the cell pellet and thoroughly mixed on ice before pipetting onto a Sigmacote-treated glass slide as previously described^[53]^. The slide was then placed into a petri dish and incubated at 37 °C for 45 min before transferring to a 6-well plate with the cell culture medium containing 3 mL media per well. BMDM were cultured overnight in MAP scaffolds with M0 media (10% v/v heat-inactivated fetal bovine serum and 1% v/v antibiotic-antimycotic in Iscove’s Modified Dulbecco’s Medium with 15 ng mL^−1^ macrophage colony-stimulating factor) before the 24-h cytokine activation. M1 activation was with 20 ng mL^−1^ LPS and IFN-γ, and M2 activation was with 20 ng mL^−1^ IL-4.

### *In vitro* Flow Cytometry Study

The cells were extracted from MAP scaffolds gels by enzymatic release of cells from the HMPs with 0.1 X TrypLE + 1mM EDTA + 1% antibiotic-antimycotic for 15 min at 37 °C. The resulting content was diluted to 10ml with PBS and filtered through a 30 µm cell strainer (CellTrics) to get a single-cell suspension and separate out the HMPs (larger than 30 µm so they won’t go through the filter). These cells were then stained with a Zombie NIR (BioLegend) solution for 15 min at room temperature and blocked with Fcr Blocking Reagent (Miltenyi Biotec) for 10 min on ice, followed by surface marker staining for 30 min on ice.

Cells were then plated in 96 well plates and viability staining was performed with Zombie-NIR (BioLegend) for dead cell exclusion. Blocking was performed with Fcr Blocking Reagent (Miltenyi Biotec). Cells were stained with the following antibody cocktail: Arg1 (eBioscience, clone A1exF5), CD11b (eBioscience, clone M1/70), CD11c (BioLegend, clone N418), CD206 (BioLegend, clone C068C2), CD86 (BioLegend clone GL-1), F4/80 (eBioscience, clone BM8), iNOS (eBioscience, clone CXNFT), MHCII (eBioscience, clone M5/114.15.2), Siglec-E (BioLegend, clone M1304A01), TLR4/MD2 (BioLegend, clone MTS510). Surface staining was conducted in FACS buffer containing 1X PBS, 0.2% w/v bovine serum albumin, 1 mM EDTA, and 0.025% w/v Proclin. Fixation, permeabilization, and intracellular staining were conducted with the eBioscience Foxp3/Transcription Factor staining buffer set. After the completion of staining, cells were resuspended in FACS buffer and run on a Cytek NL-3000 Flow Cytometer. Data were un-mixed using SpectroFlo software and analyzed using FlowJo 8 (TreeStar) flow cytometry data analysis software. The relative abundance of the macrophage population and subpopulations was determined following the gating strategy (Figure S5).

### ELISA

Cell culture medium was collected from MAP scaffolds confined BMDMs after 24 h of cytokine stimulation. To remove cell debris, the cell culture medium was centrifuged, and the supernatant was collected and stored at −80 °C until further processing. All ELISA kits were purchased from Thermo Fisher Scientific, and the tests were performed according to the manufacturer’s protocol.

### Photothrombotic Stroke

Animal procedures were performed in accordance with the US National Institutes of Health Animal Protection Guidelines and approved by the Chancellor’s Animal Research Committee as well as the Duke Office of Environment Health and Safety under the protocol number.

C57BL/6 male mice of 8–12 weeks (Jackson Laboratories), were used in the study. A photothrombotic (PT) was performed as previously described. Animals were put under isoflurane anesthesia (3% induction and 1.5% maintenance at 1 L min^−1^ USP grade O2). and placed on stereotactic apparatus. After application of eye ointment and sterilization of mice head, a midline incision was done to expose skull and connective tissue were removed. Laser was positioned 1.5 mm lateral from the bregma, and mouse was interperitoneally (IP) injected with photosensitive dye rose Bengal (Sigma; 10 mg mL^−1^ in PBS) at 10 µL g^−1^ mouse bodyweight. After 7 min, the brain was illuminated at 42 mW through intact skull for 13 min. A burr hole was drilled through the skull at 1.5 mm medial/lateral, same as laser position. The incision was then closed using Vetbond (3M, St. Paul, MN). Mice were placed in a clean cage on heat pad and monitored until awake and alert.

### Stereotactic MAP Injection

Five days post-stroke, mice were treated with 4.5 µL of MAP (M), MAP + 100% Bound PSA (M + PSA_b_), MAP + 50% Bound PSA + 50% Soluble PSA (M + PSA_b/s_), or MAP + 100% Soluble PSA (M + PSA_s_) via stereotactic injection into the stroke cavity. Mice were randomly assigned to experimental conditions within each cage to control for potential cage-specific confounding factors. For treatment injections, mice were first anesthetized with isoflurane at 3% and then transferred and positioned on a stereotactic stage for isoflurane maintenance at 1.5%. Following alcohol and iodine washes (3x), the skin covering the skull was re-opened, exposing the location of the photo-thrombotic stroke. A MAP treatment (4.5 µL) consisting of packed HMP (with or without bound PSA) and 0.25 mM HA-Tet solution at a volume ratio of 5:1 was injected directly into the stroke cavity through a flat 30-gauge needle attached to a 25 µL Hamilton syringe (Hamilton, Reno, NV). A syringe pump was used to deliver treatments at an infusion rate of 1 µL min^−1^at the same coordinates as the PT stroke laser irradiation, 1.5 mm lateral from bregma, and 0.75 mm ventral from the skull.

Prior to injection, the syringe was lowered 0.75 mm into the infarct site. Five minutes after completion of the injection, the syringe was slowly lifted out of the infarct site. The incision was then closed using Vetbond (3M, St. Paul, MN). Mice were placed in a clean cage on heat pad and monitored until awake and alert.

### *In vivo* flow Cytometry Study

At the terminal endpoint (day 7), mice were anesthetized with isoflurane and perfused with at least 10 mL of ice cold 1x PBS. Following perfusion, brains were extracted, and the ipsilateral hemisphere was isolated for further tissue processing and analysis. Tissues were disassociated enzymatically in a solution of 1 mg/mL collagenase (Sigma-Aldrich) and 1 U/mL DNase (Worthington Biochemical) in RPMI-1640 with L-glutamine (Corning) supplemented with 2% v/v fetal bovine serum (R&D Systems), 1% v/v penicillin/streptomycin (Thermo Fisher), and 10 mM HEPES (Gibco) at 37 °C with mechanical dissociation by shaking with 0.25 inch ceramic beads (MP Biomedicals) for 25 minutes. Dissociated tissues were passed through 70 µm cell strainers, and a 40% Percoll (Cytiva) density gradient was used to separate cells from myelin debris. Cells were then plated in 96 well plates and viability staining was performed with Zombie-NIR (BioLegend) for dead cell exclusion. Blocking was performed with Fcr Blocking Reagent (Miltenyi Biotec). Cells were stained with the following antibody cocktail: Arg1 (Thermo Fisher, clone A1exF5), B220 (BioLegend, clone RA3-6B2), CD11b (Thermo Fisher, clone M1/70), CD11c (Thermo Fisher, clone N418), CD206 (BioLegend, clone C068C2), CD3 (BioLegend, clone 17A2), CD45 (Thermo Fisher, clone 30-F11), CD64 (BD Biosciences, clone X54-5/7.1), CD86 (BioLegend, clone GL-1), F4/80 (BioLegend, clone BM8), iNOS (Thermo Fisher, clone CXNFT), Ly6C (BioLegend, clone HK1.4), Ly6G (Thermo Fisher, clone 1A8), MHCII (Thermo Fisher, clone M5/114.15.2). Surface staining was conducted in FACS buffer containing 1x PBS with 0.5% w/v bovine serum albumin (R&D Systems) and 2mM EDTA (Santa Cruz Biotechnology). Fixation, permeabilization, and intracellular staining were conducted with the eBioscience Foxp3/Transcription Factor staining buffer set. Finally, cells were resuspended in FACS, and flow cytometry was run on a Cytek NL-3000 Flow Cytometer.

### Tissue Processing for Immunostaining

At the terminal endpoint (day 15), mice were anesthetized with isoflurane and perfused with at least 10 mL of ice cold 1x PBS followed by ice cold 4% (w/v) paraformaldehyde (PFA, Electron Microscopy Sciences). Following perfusion, brains were extracted and placed in a 4% PFA solution for 4 h at 4 °C. Brains were then washed three times with 1X PBS, after which they were placed into a 30% sucrose solution prepared in 1X PBS at 4 °C. Brains were cryosectioned into 30 µm slices, collected onto glass slides, and preserved at −80 °C until immunostaining.

### Immunostaining

In preparation for immunofluorescent (IF) staining, slides were warmed to room temperature. Slides were then washed in 1x TBS (Sigma, D8537) for 5 min three times. A blocking buffer, comprised of 1x TBS, 0.3% Triton-X, and 10% normal donkey serum (NDS), was used to block tissue samples at room temperature for 1–2 h. Primary antibodies, prepared in blocking buffer, were then left to incubate with tissue overnight at 4 °C. The next day, slides were washed in 1x TBS + 0.3% Triton-X for 10 min three times. Secondary antibodies and DAPI (1:1000), prepared in blocking buffer, were then left to incubate with tissue for 2 h at room temperature, protected from light. Slides were then washed quickly with 1x TBS, followed by three additional 10 min 1x TBS washes. Tissue was left to dry at room temperature for ≈1–2 h. To dehydrate tissue sections, slides were placed in a series of ethanol baths: 50%, 50%, 70%, 70%, 90%, 90%, 95%, 95%, 100%, and 100%. Slides were left in each bath for ≈1 min, except for the final 100% ethanol bath, during which slides were left to incubate for at least 3 min. Following dehydration, tissue was then defatted in two xylene baths: 1 min. in the first bath and at least 5 min in the second bath. Slides were then mounted in DPX and left to dry overnight at room temperature. *Primary Antibodies*: 1:250 Iba1 (Wako Chemicals, 019– 19741), 1:250 Tmem119 (Synaptic Systems, 400 004). *Secondary Antibodies*: 1:500 Alexa Fluor 555 (Guinea Pig, Invitrogen, A-21450), 1:500 Alexa Fluor 555 (Rb, Invitrogen, A- 31572).

### Imaging and Image Analysis

All fluorescence imaging was performed using a Nikon Ti Eclipse microscope equipped with a C2 laser with a 20× air objective at a 4 µm step-size. Image processing and analysis were conducted in Fiji/ImageJ. Images are presented as maximum intensity projections (MIPs).

For quantification of Iba1+ area, images were despeckled, thresholded, and analyzed within appropriate regions of interest (ROIs). Signal normalization was carried out by dividing the Iba1+ area by the void space, identified via thresholding of a DAPI channel subjected to Gaussian blur (σ = 8). To determine the proportion of microglia in Iba1+ cells, the Iba1+ area was used as a ROI mask and applied to the Tmem119 channel, enabling quantification of Tmem119+ signal restricted to Iba1+ regions.

### Statistical Analysis

All *in vitro* BMDM experiments were performed in biological triplicate, with each replicate comprising BMDMs isolated and cultured from a single mouse. Flow cytometric screens of endotoxin containing PSA or sialic-acid treatments and the BMDM-in-MAP encapsulation study were analyzed by one-way ANOVA; when the F-test was significant (p < 0.05), pairwise differences were resolved with Tukey’s HSD. Siglec-E qPCR validation was assessed with a two-way ANOVA (fixed factors: primer set and cell type) using a single pooled error term. The primer main effect was nonsignificant, but a significant primer x cell-type interaction (p < 0.05) prompted Sidak-adjusted comparisons that were limited to cell-type contrasts within each primer set (α = 0.05). Material characterization (rheometry, enzymatic degradation, and void-space modelling) employed three to five independently cast MAP scaffolds, each treated as an independent technical replicate for statistical calculations. For *in vivo* immunofluorescence quantification, the sample size was determined *a priori* with G*Power v3.1.9.6 (two-tailed, unpaired Student’s t-test, α = 0.05, power = 0.80). Pilot data yielded an effect size of Cohen’s d = 1.77, indicating that 6 animals per group were required. For each animal, values from two brain sections were averaged, and this mean served as a single biological replicate. Results are plotted as floating-bar graphs (min–max) with individual animal means overlaid. Homogeneity of variance was confirmed (Levene’s test, p > 0.05), and group differences were evaluated with a two-tailed, unpaired Student’s t-test using the pooled-variance estimate (α = 0.05). For the *in vivo* brain flow cytometry study, statistics were performed in two stages. First, a three-way ANOVA (fixed factors: cell type, treatment, and marker) was first used to probe main effects and interactions. For significant interactions (p < 0.05), the data set was stratified by the relevant factor and re-examined with a one-way ANOVA followed by Tukey’s post-hoc test to compare treatment groups within each stratum. An *a priori* power analysis in G*Power (two-tailed α = 0.05, power = 0.80, Cohen’s d = 1.97 for immune cell count and Cohen’s d = 2.63 for microglia percentage derived from pilot work) indicated a target of 4-6 animals per group; logistical limits in animal-housing capacity and study budget restricted enrolment to 4 animals per group. For all experiments, significance was indicated by *P < 0.05, **P < 0.01, ***P < 0.001, and ****P < 0.0001. Statistical analyses for *in vitro* experiments, material characterizations, and *in vivo* immunofluorescence imaging quantification were performed with GraphPad Prism v10.5.1. For *in vivo* brain flow cytometry data analysis, the three-way ANOVA test was performed in JMP v18.2.1, and the following one-way ANOVA was performed with GraphPad Prism v10.5.1.

## Acknowledgements

The authors would like to thank Dr. Juhi Samal for the acquisition of brain glycan data that inspired this work. The authors would like to thank Alejandra Suarez-Arnedo for her time in teaching the BMDM extraction and *in vitro* flow cytometry technique used in this work. The authors would like to thank Yasha Saxena in developing the computational algorithm used to reconstruct confocal images for LOVAMAP analysis. The authors would like to thank Dr. Lindsay Riley for her time in teaching the LOVAMAP pipeline. The authors would like to thank Dr. Andrea Jones for kindly helping with outlining and editing the manuscript. Lastly, we would like to acknowledge our funding from the National Institutes of Health through R01NS079691 (TS) and R01NS112940 (TS).

## Conflicts of Interest

Y.O. and T.S. are co-inventors on a provisional patent application (U.S. Provisional Patent Application No. 63/784,550), filed by Duke University, entitled ‘GLYCAN ACTIVE MICROPOROUS ANNEALED PARTICLE GEL SYSTEM AND METHOD THEREOF,’ which relates to the polysialic acid-functionalized MAP technology described in this manuscript. The other authors declare no competing interests.

Received: ((will be filled in by the editorial staff))

Revised: ((will be filled in by the editorial staff))

Published online: ((will be filled in by the editorial staff))

## Supporting Information

**Figure S1.**
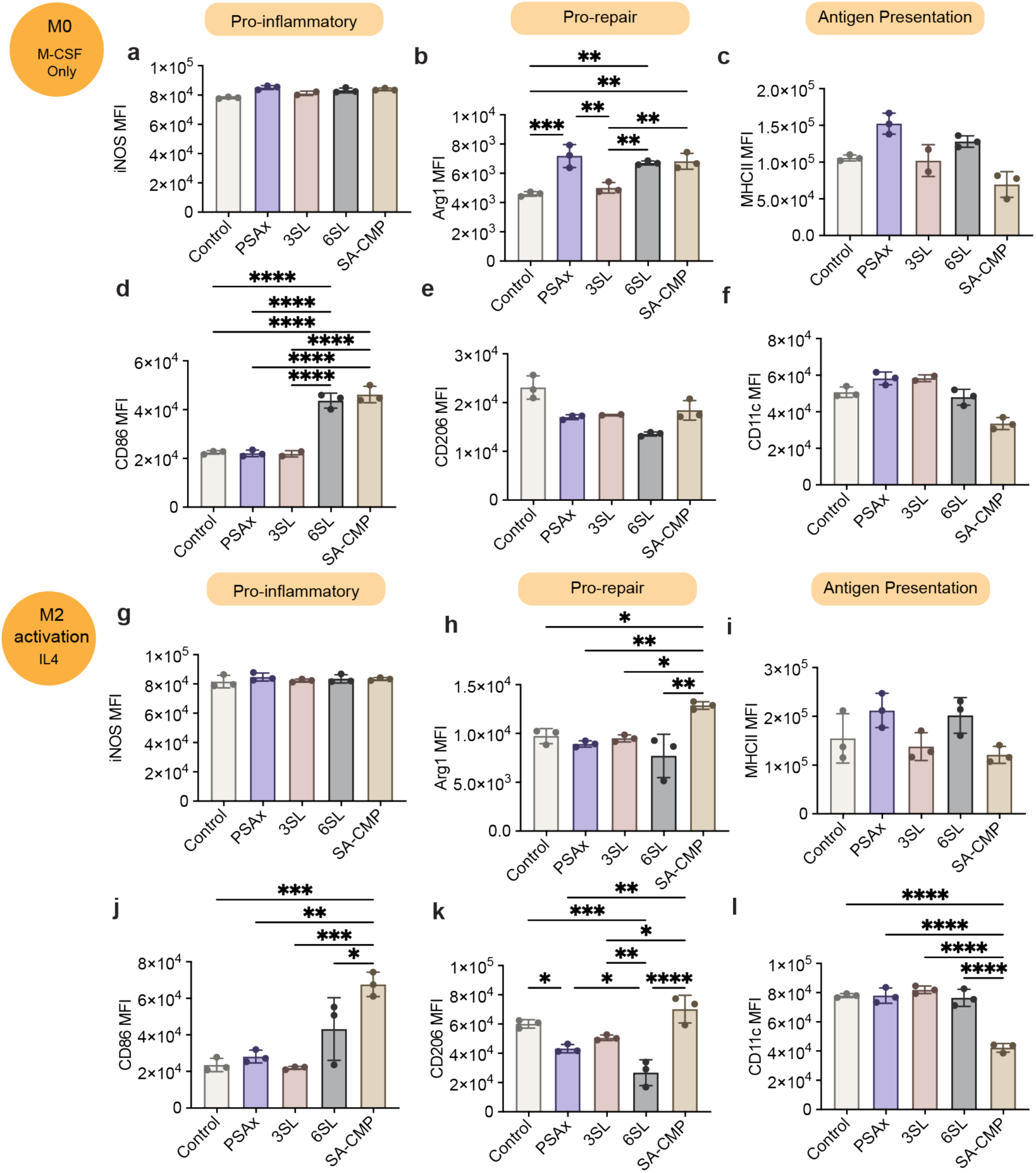
Sialic acid derivative screening under M0 and M2 macrophage activation. **a–f**, Macrophage marker expression (MFI) under M0 baseline conditions. Significant increases in Arg1 were observed with PSAx, 6’SL, and CMP-SA treatments (**b**). Notably, 6’SL and CMP-SA significantly increased CD86 expression compared to untreated M0 control (**d**). **g–l**, Marker expression under M2 conditions (IL-4 activation). CMP-SA significantly elevated Arg1 (**h**) and CD86 (**j**) while decreasing CD11c (**l**). PSAx and 6’SL significantly reduced CD206 (**k**). Data presented as mean ± SEM, n = 3 biological replicates; *p<0.05, **p<0.01, ***p<0.001, ****p<0.0001, determined by one-way ANOVA with Tukey’s post-hoc test.

**Figure S2.**
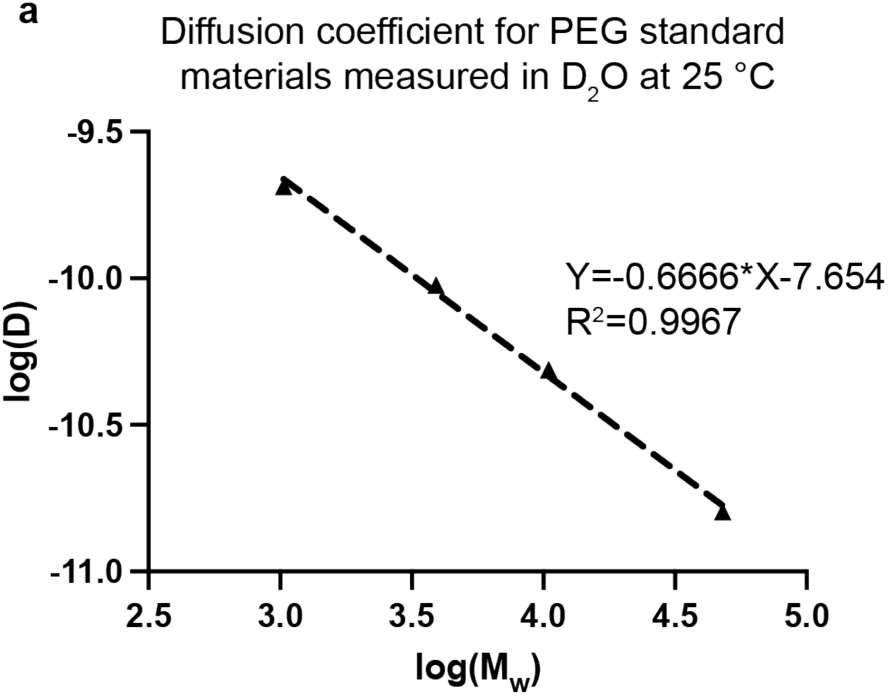
DOSY spectroscopy standard curve. **a**, DOSY standard curve correlating the diffusion coefficient (log D) and molecular weight (log MW), established using PEG standards measured in D₂O at 25°C. This calibration curve enabled molecular weight determination for various PSA derivatives in Figure 3.

**Figure S3.**
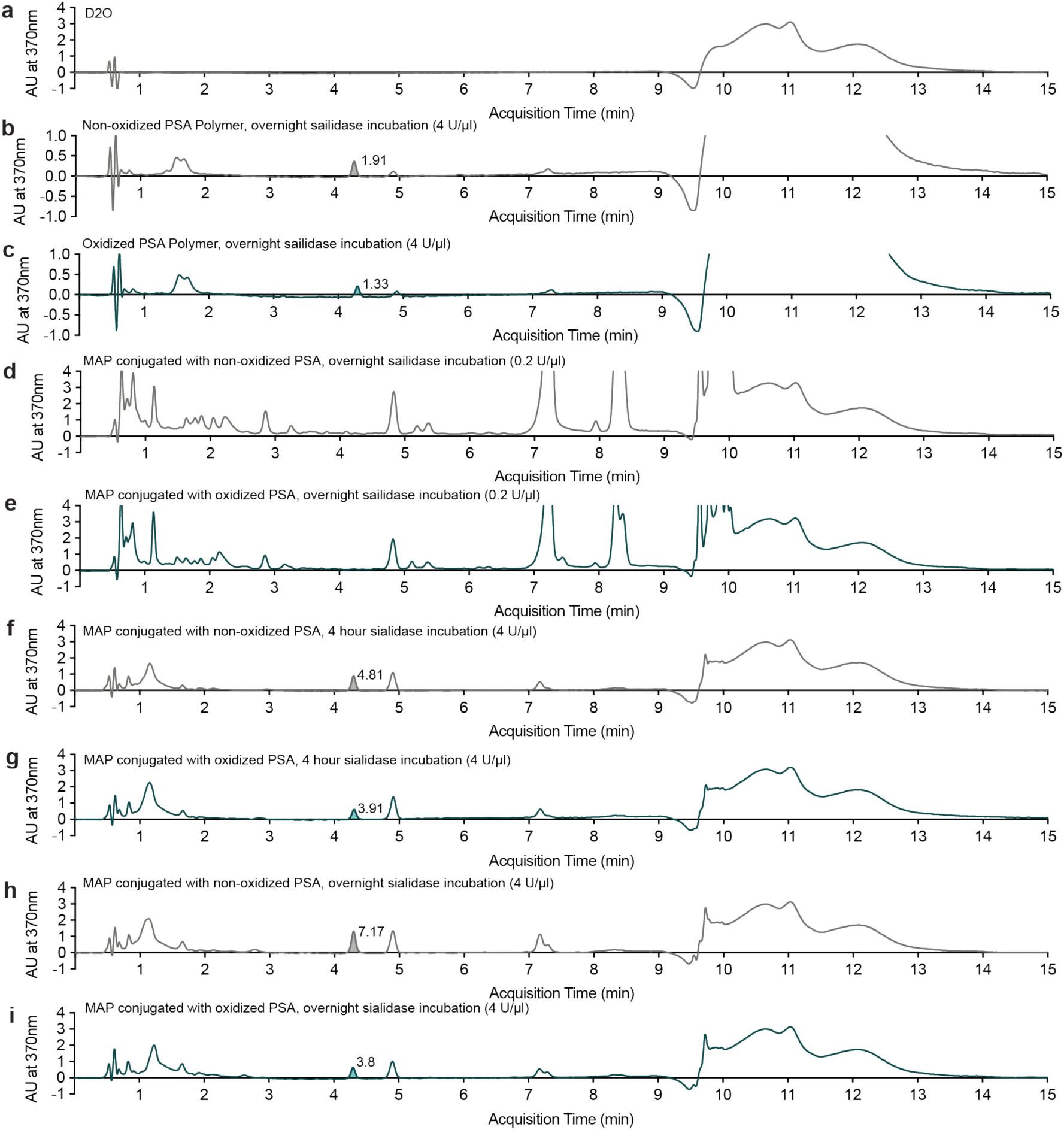
HPLC chromatograms of PSA degradation under various conditions. Representative full-range (0–15 min) HPLC chromatograms for: **a**, Water control; **b-c**, Non-oxidized and oxidized PSA polymers (not conjugated to MAP scaffolds) incubated overnight at high sialidase concentration (4 U/µL); **d-e**, MAP scaffolds conjugated with non-oxidized and oxidized PSA incubated overnight at physiological sialidase concentration (0.2 U/µL); **f-g**, MAP scaffolds conjugated with non-oxidized and oxidized PSA incubated for 4 hours at high sialidase concentration (4 U/µL); **h-i**, MAP scaffolds conjugated with non-oxidized and oxidized PSA incubated overnight at high sialidase concentration (4 U/µL). Peaks at ∼4.3 min indicate enzymatically released Neu5Ac following PSA degradation with DMB labeling.

**Figure S4.**
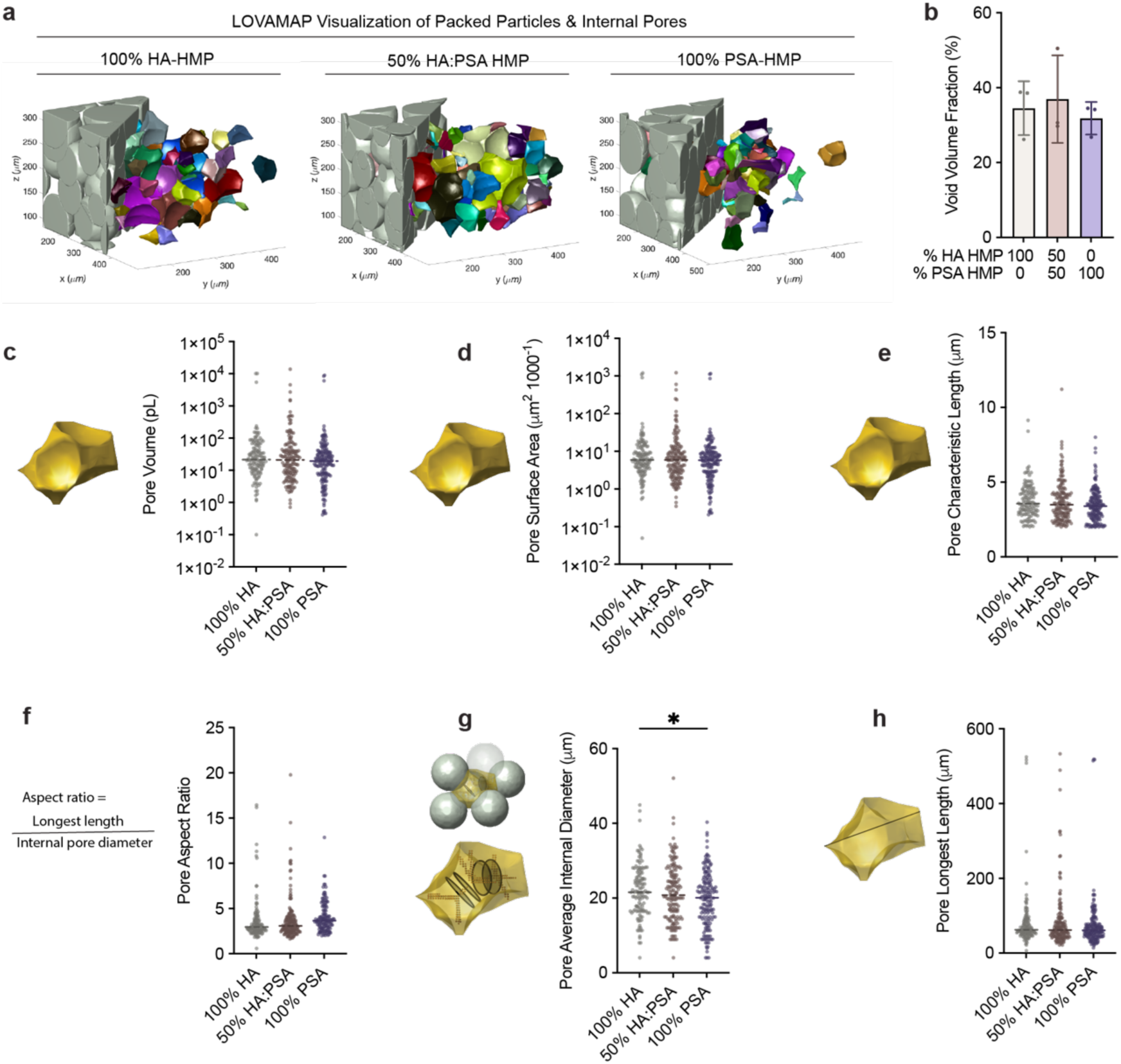
Computational LOVAMAP analysis of void space of MAP scaffolds. **a**, LOVAMAP outputs of MAP scaffolds. HMPs are grey and 3D-pores are colored. **b**, Void fraction in MAP scaffolds. Void fraction = (Total volume – Microgel volume) / Total volume. Additional LOVAMAP outputs for the distribution of pore volume (**c**), surface area (**d**), characteristic length (**e**), aspect ratio (**f**), average internal diameter (**g**), and longest length (**h**). Data presented as mean ± SEM (n=3 independently casted scaffolds per formulation). One-way ANOVA with Tukey’s post hoc test: *p < 0.05.

**Figure S5.**
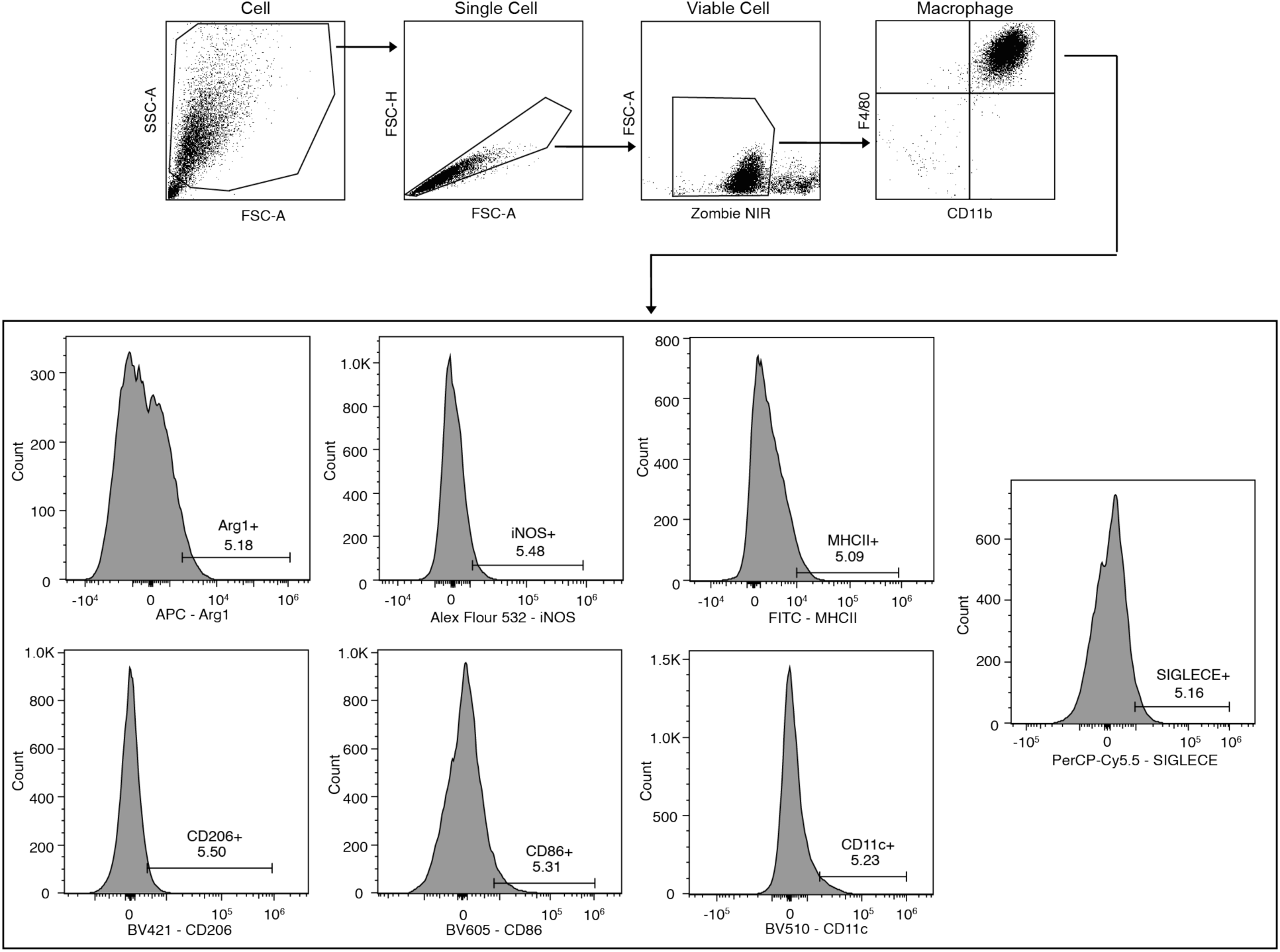
Gating strategy for flow cytometry analysis on bone marrow derived macrophages with 10 markers.

**Figure S6.**
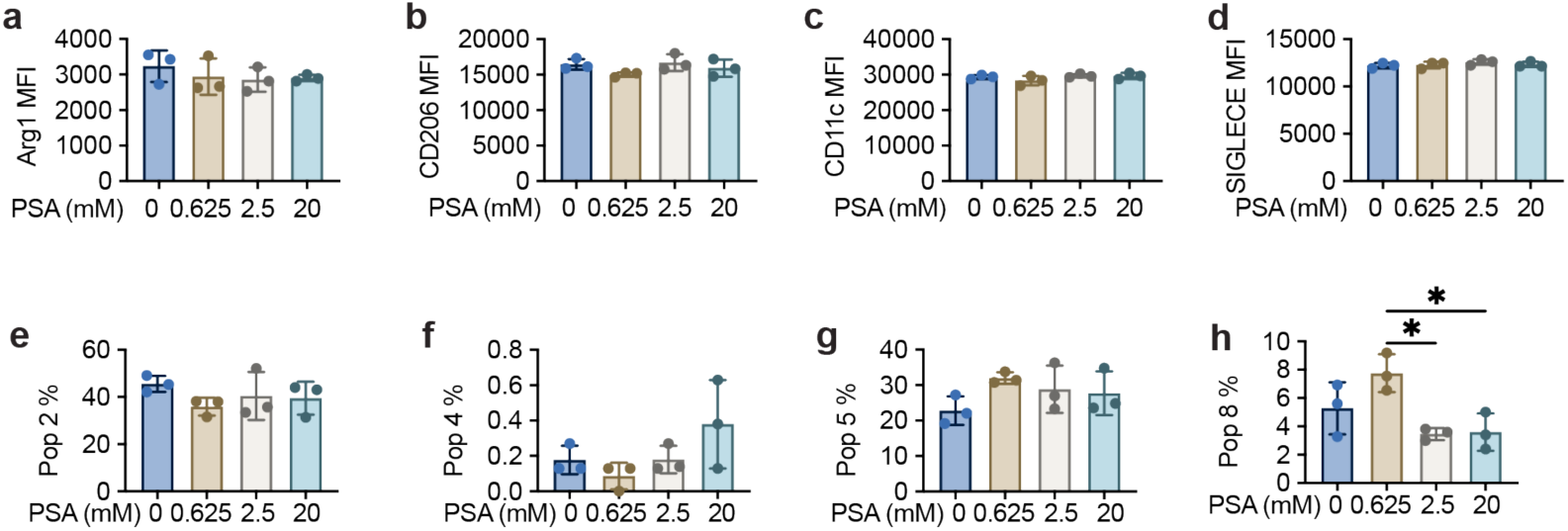
Additional phenotypic marker expression in BMDMs cultured within PSA-MAP scaffolds. Mean fluorescence intensities (MFI) of additional BMDM phenotypic markers following encapsulation in MAP scaffolds with varying PSA concentrations: **a**, Arg1; **b**, CD206; **c**, CD11c; **d**, SIGLEC-E. **e-h**, Relative frequencies of additional cell populations (Populations 2, 4, 5, and 8) from high-dimensional clustering analysis (UMAP) across PSA concentrations, revealing subtle changes in macrophage phenotypic states. Data represent mean ± SEM (n = 3 biological replicates; each replicate represents cells isolated from a distinct mouse). One-way ANOVA with Tukey’s post-hoc test: *p < 0.05.

**Figure S7.**
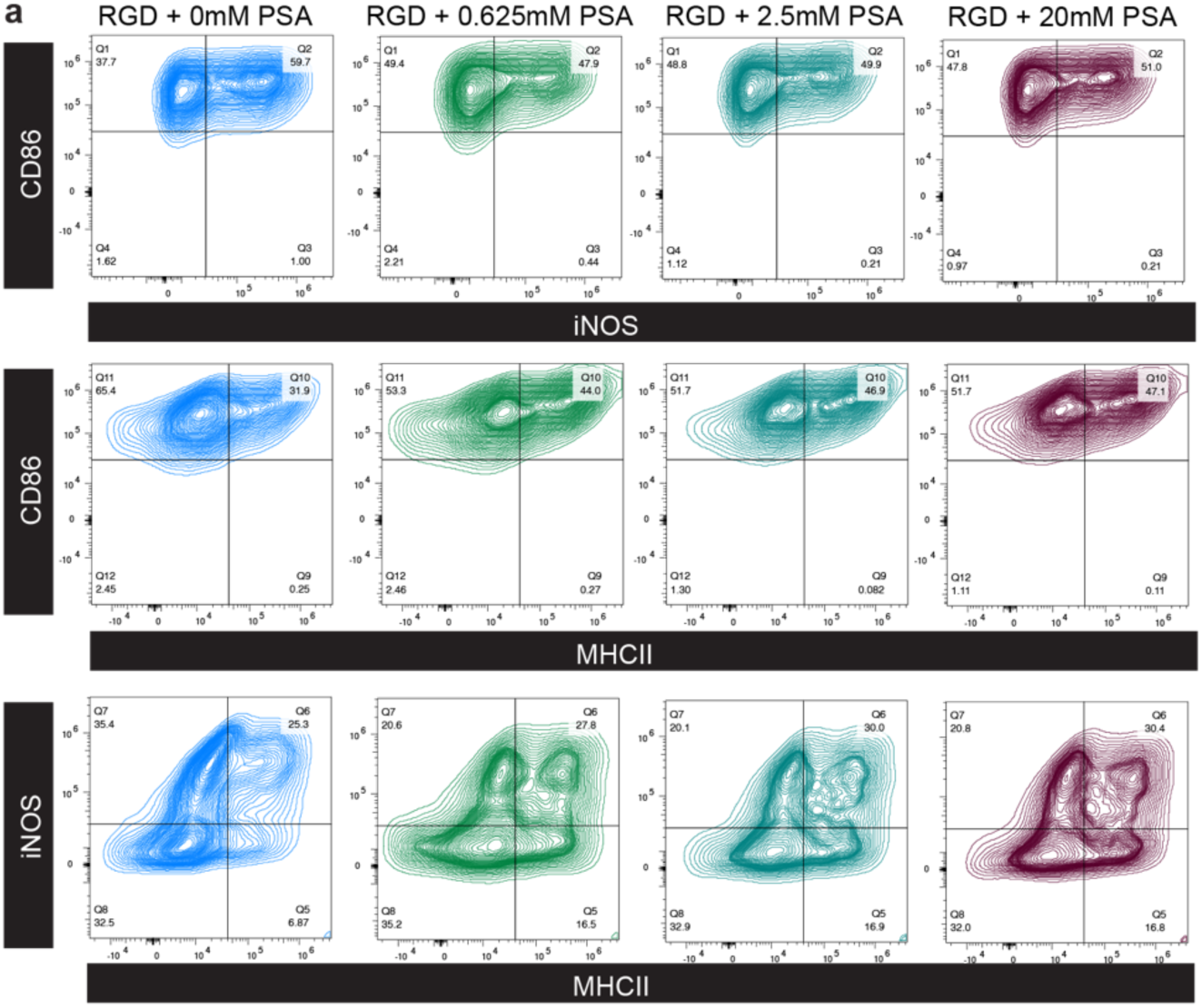
Flow cytometry double-marker contour plots for BMDMs. Representative contour plots for double-marker gating analysis of BMDMs encapsulated within MAP scaffolds at different PSA concentrations. Columns represent PSA concentrations (0, 0.625, 2.5, and 20 mM, left to right). Rows represent gating strategies,

**Figure S8.**
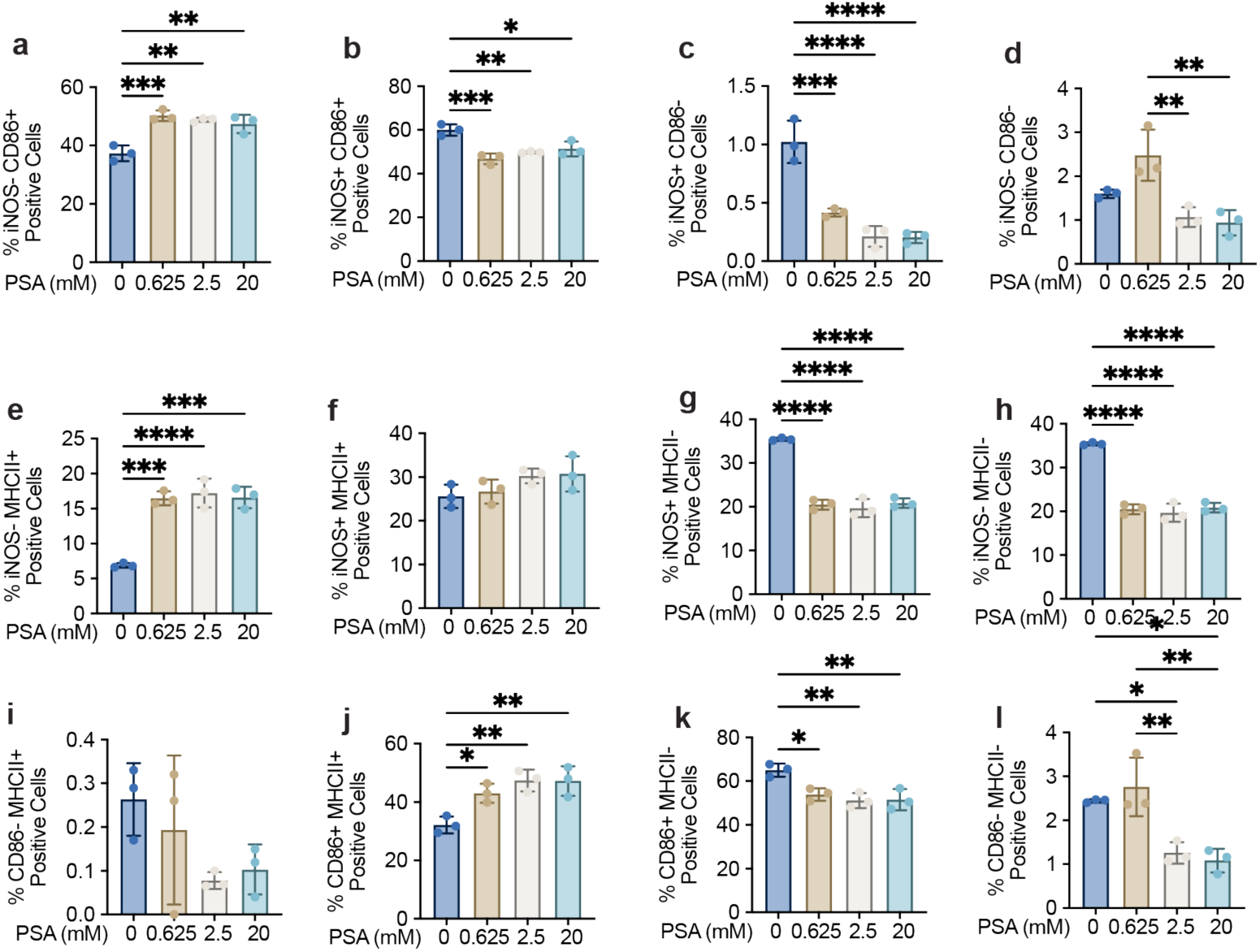
Quantification of double-marker positive BMDMs subpopulations. Quantitative analyses from double-marker gating shown in Supplemental Figure S4: **a-d**, iNOS vs. CD86 subpopulations (iNOS⁻CD86⁺, iNOS⁺CD86⁺, iNOS⁺CD86⁻, and iNOS⁻CD86⁻). **e-h**, iNOS vs. MHCII subpopulations (iNOS⁻MHCII⁺, iNOS⁺MHCII⁺, iNOS⁺MHCII⁻, and iNOS⁻MHCII⁻). **i-l**, CD86 vs. MHCII subpopulations (CD86⁻MHCII⁺, CD86⁺MHCII⁺, CD86⁺MHCII⁻, and CD86⁻MHCII⁻). Percentages illustrate nuanced phenotypic shifts in macrophages with PSA treatment. Data represent mean ± SEM (n = 3 biological replicates; each replicate represents cells isolated from a distinct mouse). One-way ANOVA with Tukey’s post-hoc test: *p < 0.05, **p < 0.01, ***p < 0.001, ****p < 0.001.

**Figure S9.**
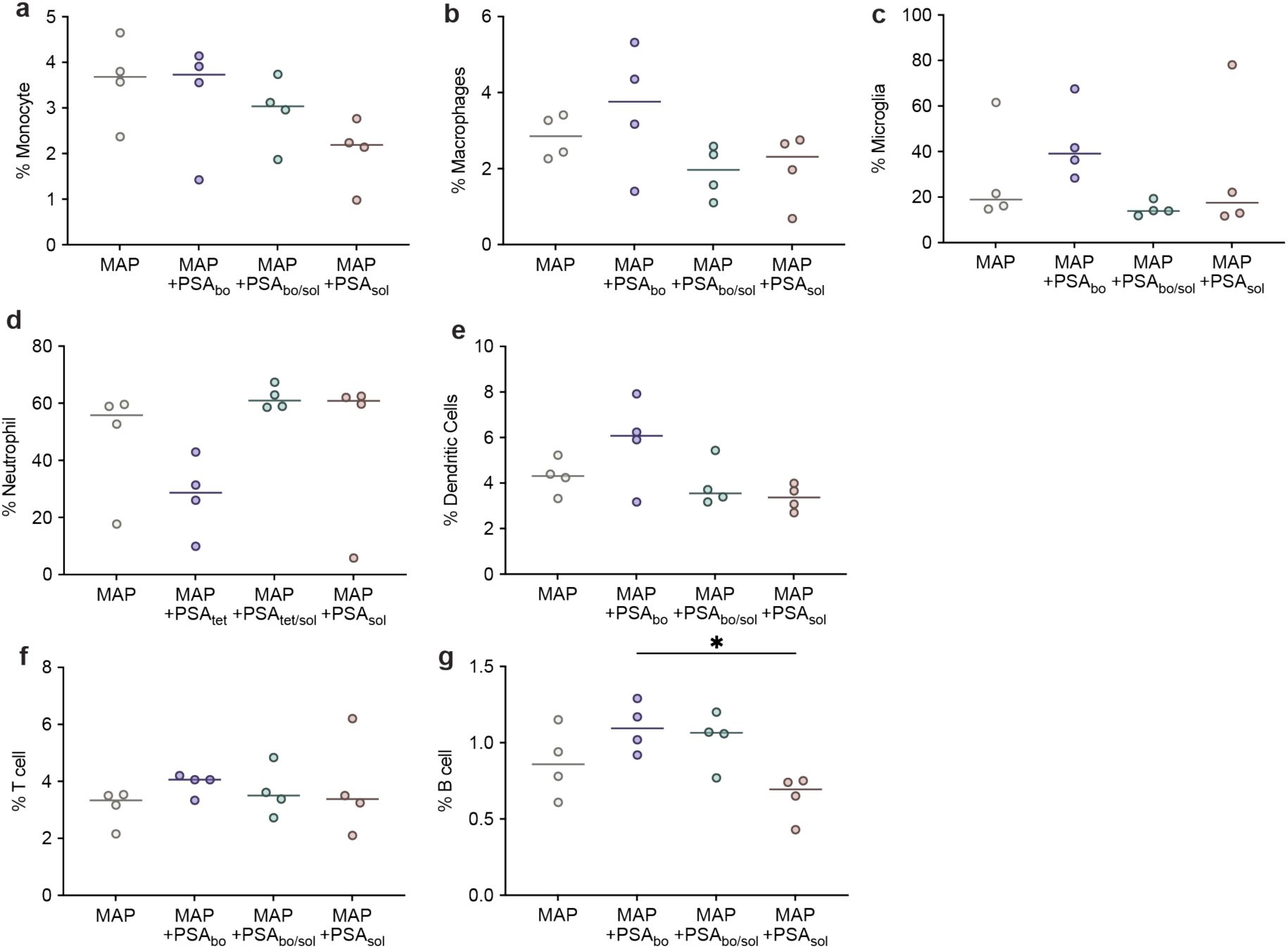
Individual data points for immune cell infiltration after MAP scaffold injection. Individual sample data points showing the percentage composition of each immune cell type (neutrophil, microglia, macrophage, monocyte, dendritic cells, B cells, and T cells) across all treatment conditions (MAP, PSAb, PSAb/s, and PSAs).

**Figure S10.**
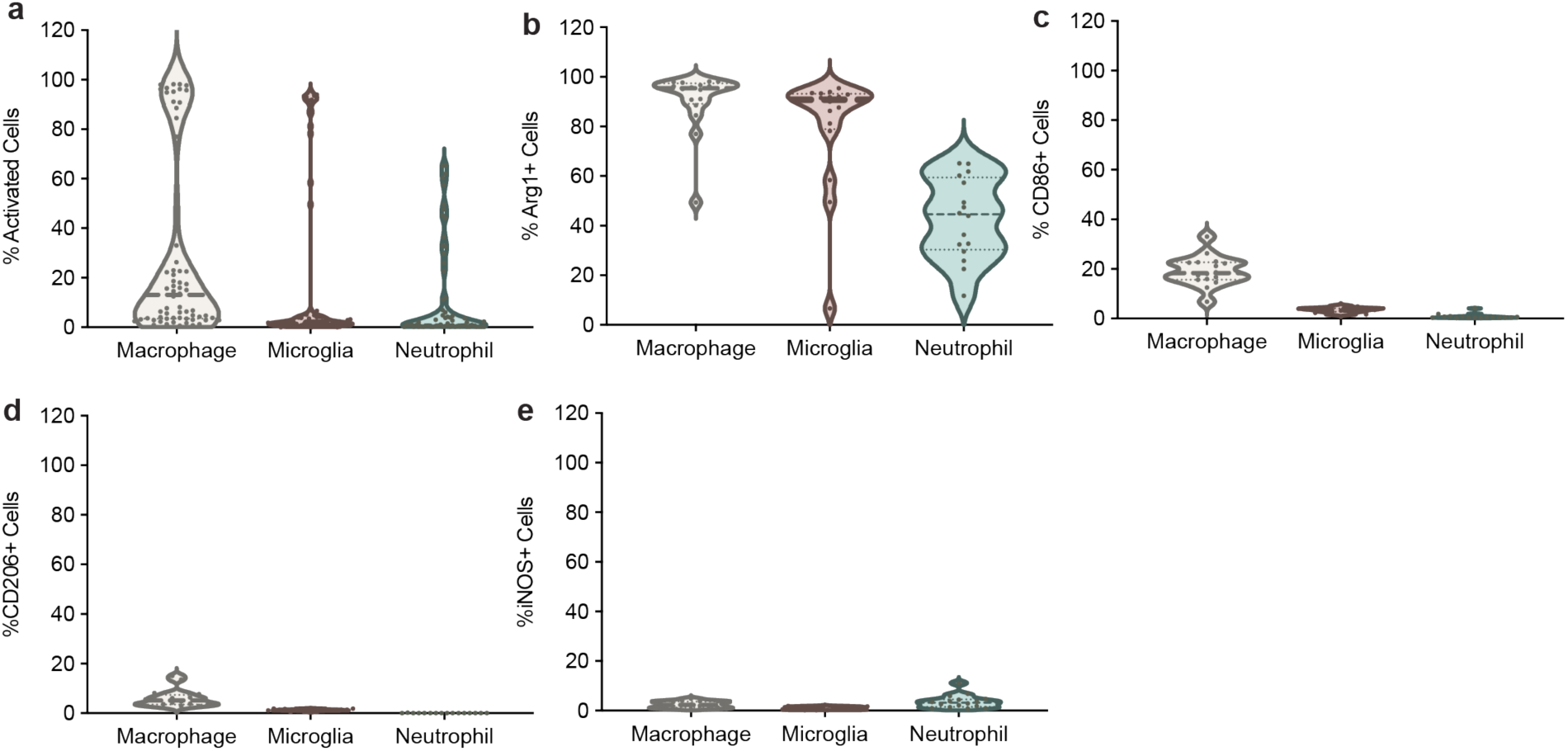
Marker expression within neutrophils, macrophages, and microglia. **a**, Percentage of activated cells (positive for Arg1, CD206, CD86, or iNOS) within each respective cell type. **b-e**, Marker-specific analyses (Arg1, CD86, CD206, and iNOS), illustrating differences in expression patterns between neutrophils, macrophages, and microglia, and highlighting distinct functional states.

**Figure S11.**
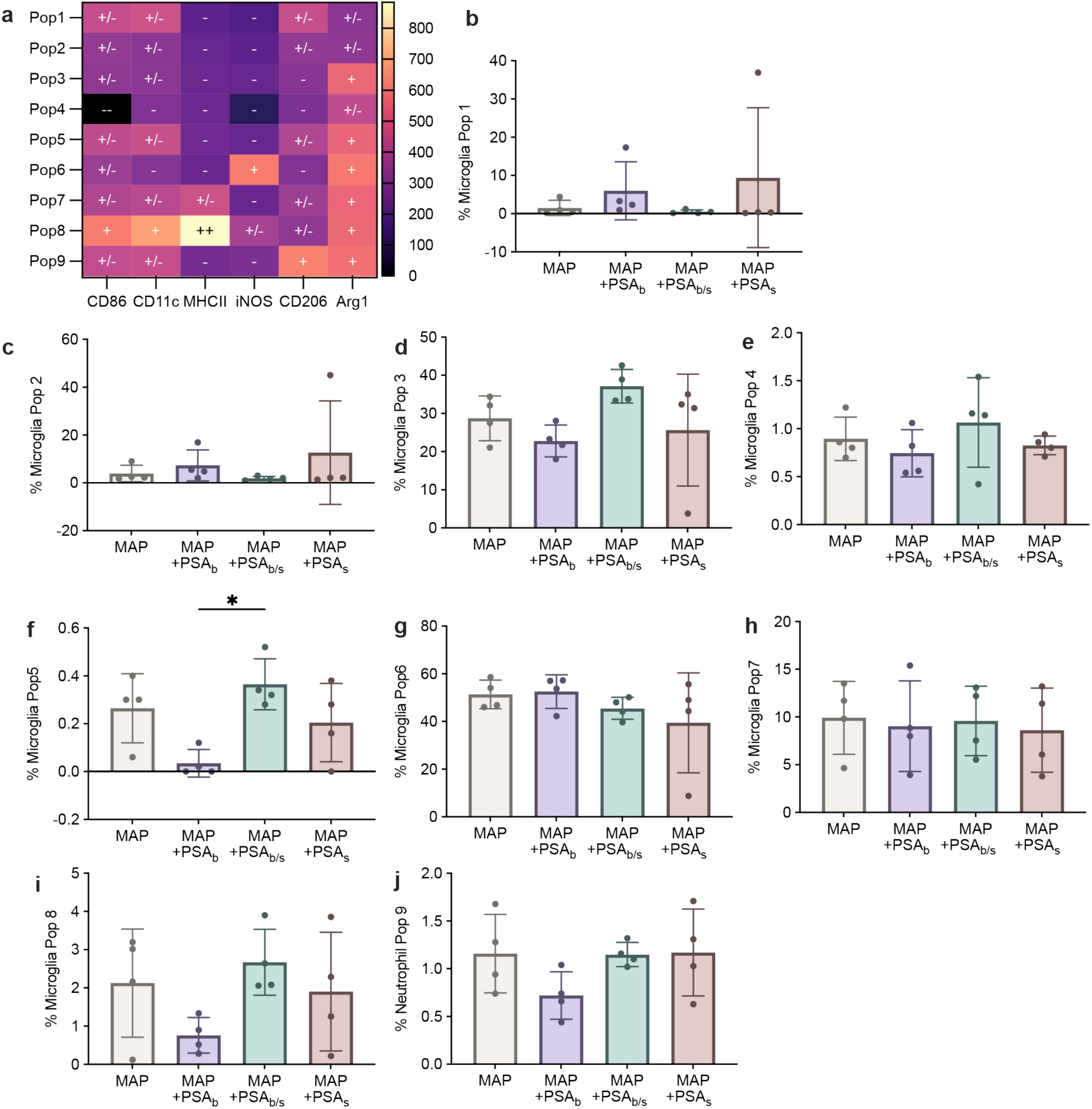
High-dimensional clustering of microglia populations (UMAP). UMAP clustering analysis identifying nine distinct microglial subpopulations. Differences among scaffold conditions (MAP, PSAb, PSAb/s, and PSAs) highlight subtle shifts in phenotypic markers, particularly Population 5 (medium CD86, Arg1, CD206 expression) and Population 8 (high CD86, CD11c, MHCII expression).

**Figure S12.**
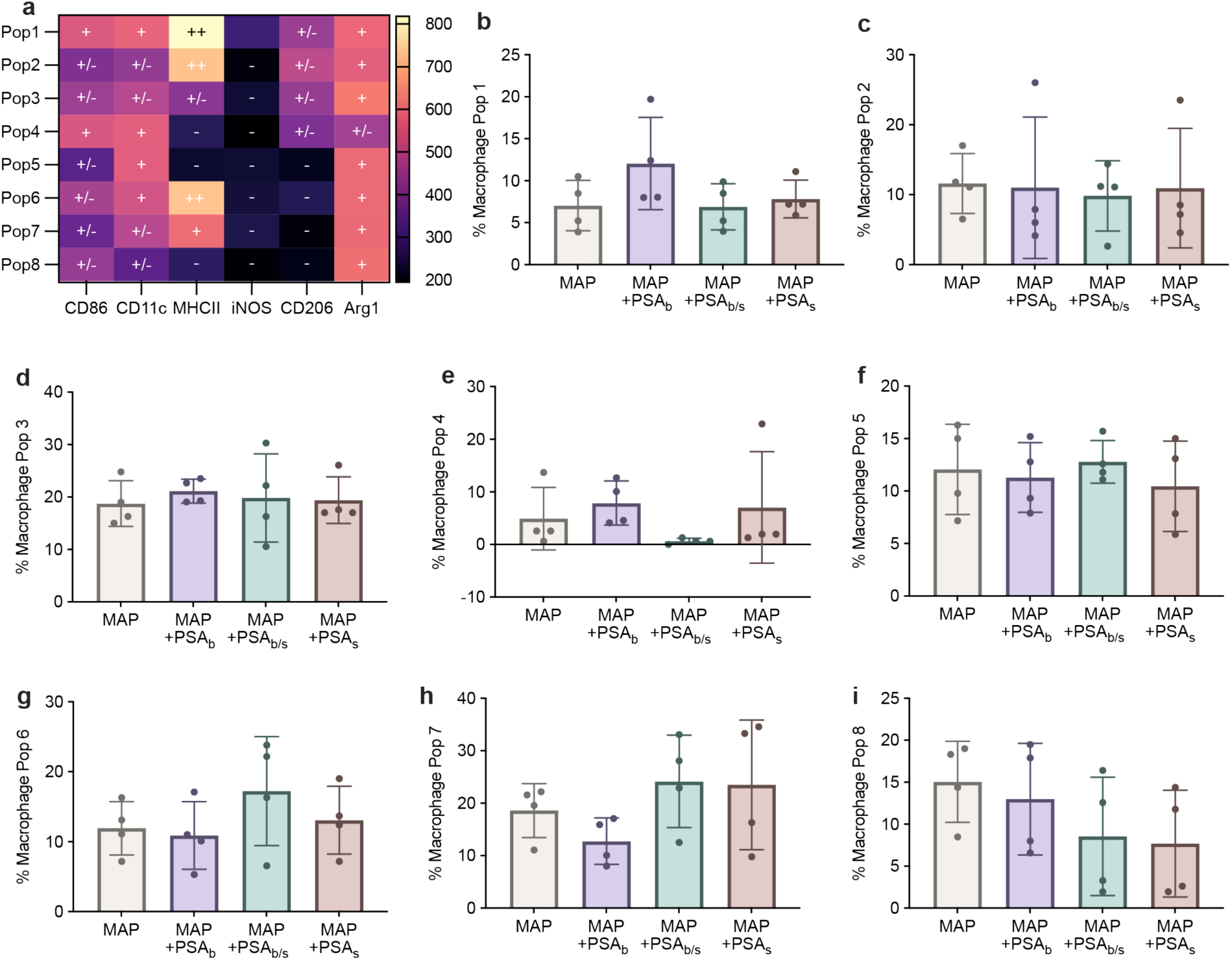
High-dimensional clustering of macrophage populations (UMAP). UMAP clustering analysis identifying distinct macrophage populations with notable phenotypic diversity. Population 1 (high MHCII, medium Arg1 and CD86) and Population 4 (medium-high CD86 and CD11c) exhibit subtle enrichment in PSA-bound scaffold conditions, suggesting nuanced polarization responses.

**Figure S13.**
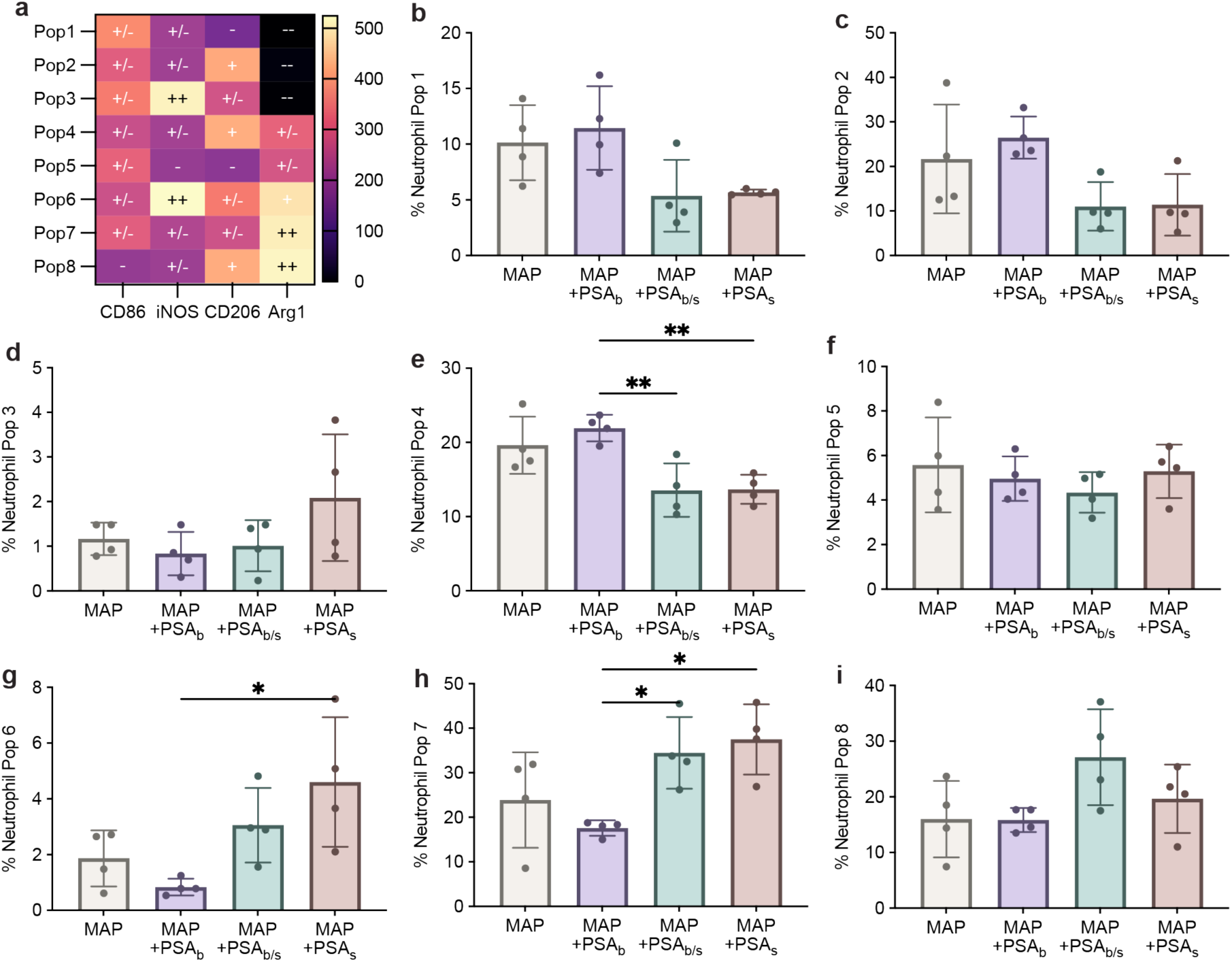
High-dimensional clustering of neutrophil populations (UMAP). UMAP clustering analysis of neutrophils (CD86, iNOS, CD206, and Arg1 expression). Notable populations include Population 4 (high CD206, medium Arg1), Population 6 (high iNOS and Arg1), and Population 7 (high Arg1, medium-high CD86), illustrating shifts toward less inflammatory phenotypes with tethered PSA scaffolds.

**Figure S14.**
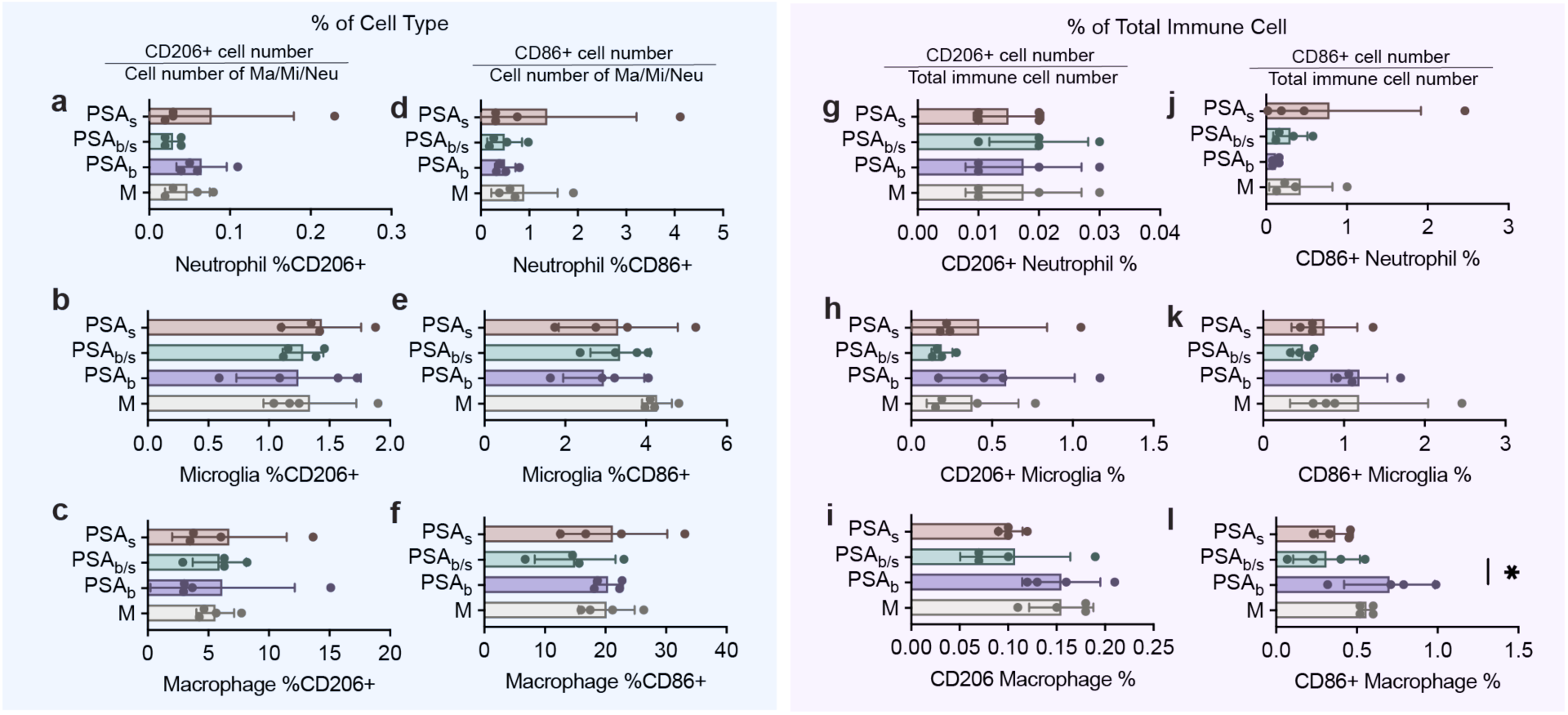
Additional marker Expression in neutrophils, macrophages, and microglia In Vivo. Additional traditional gating analyses (CD86 and CD206 expression) in neutrophils, macrophages, and microglia at day 7 post-stroke across all scaffold conditions (MAP, PSAb, PSAb/s, and PSAs). Results support main findings presented in Figure 7, showing limited significant differences in these specific markers compared to Arg1 and iNOS.

**Figure S15.**
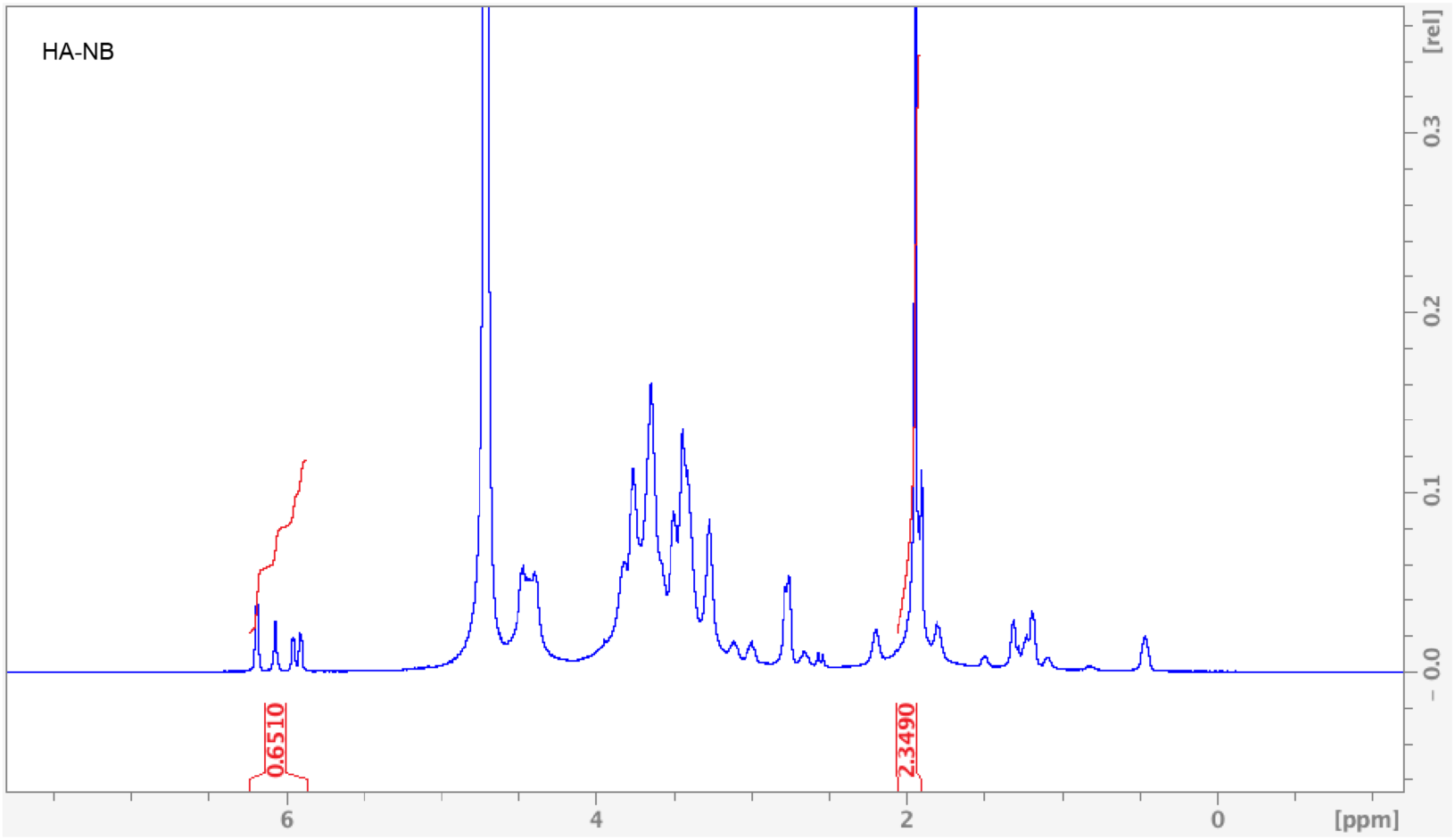
^1^H NMR analysis of HA-NB. ^1^H NMR signals in δ6.33 and δ6.02 (vinyl protons, endo), and δ6.26 and δ6.23 ppm (vinyl protons, exo) represent pendant norbornenes in D_2_O.

**Figure S16.**
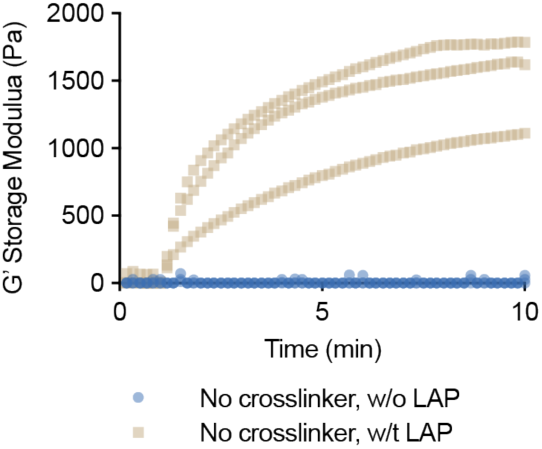
Time-sweep rheological analysis demonstrating unexpected gelation upon UV exposure (initiated at 1 min) of solutions containing HA-norbornene (HA-NB) and polysialic acid (PSA) without additional thiol-based crosslinkers, either in the presence or absence of LAP photoinitiator. Rapid increase in storage modulus (G’) in the presence of LAP indicates unintended crosslinking between PSA and HA-NB under photoinitiated conditions (n=3 independent samples per condition).

**Figure S17.**
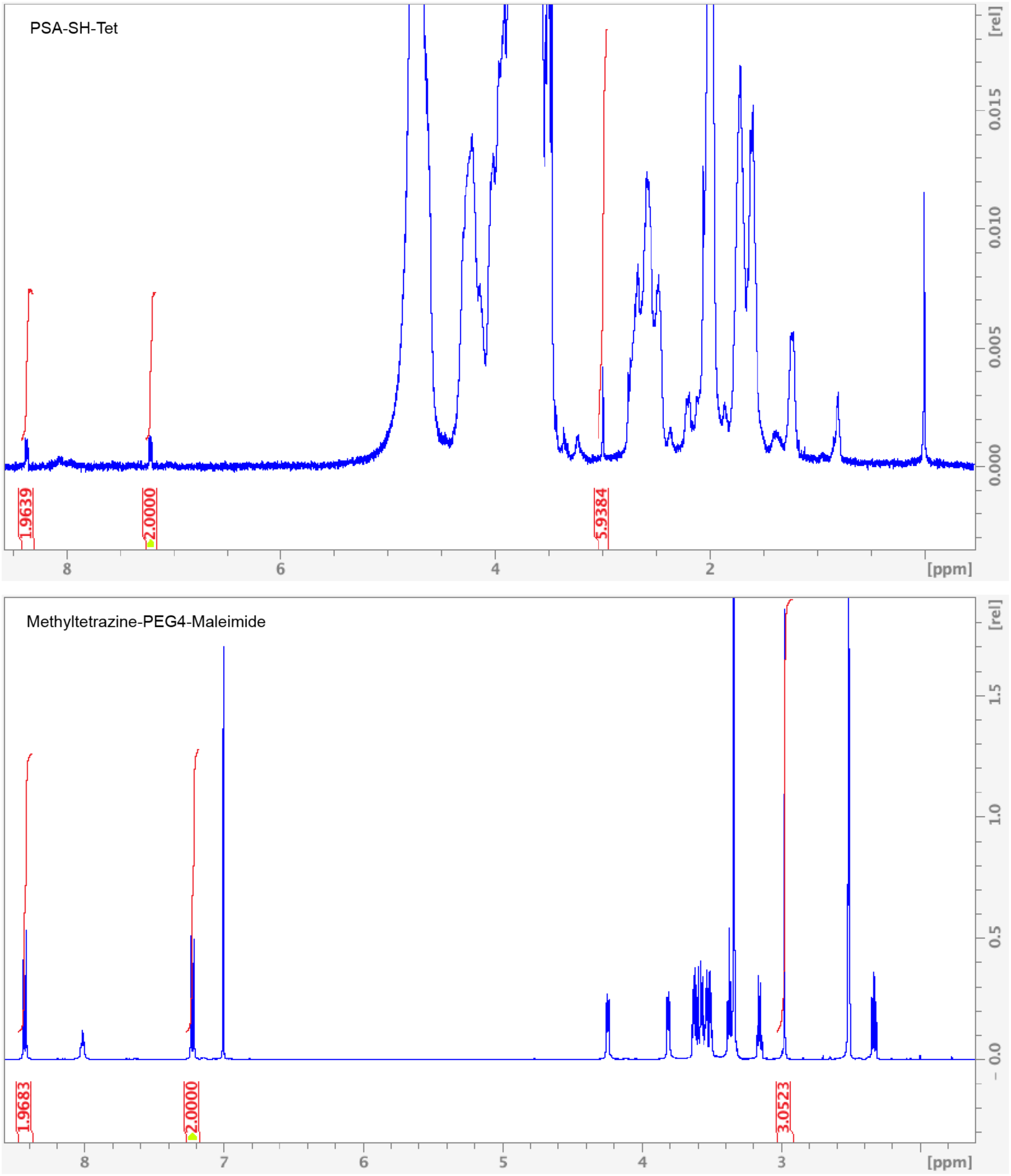
^1^H NMR analysis of PSA-Tet and methyltetrazine-PEG4-maleimide. ^1^H NMR signals at δ8.35 (2H) and δ7.20 (2H) (aromatic protons) represent tetrazines in D_2_O.

**Figure S18.**
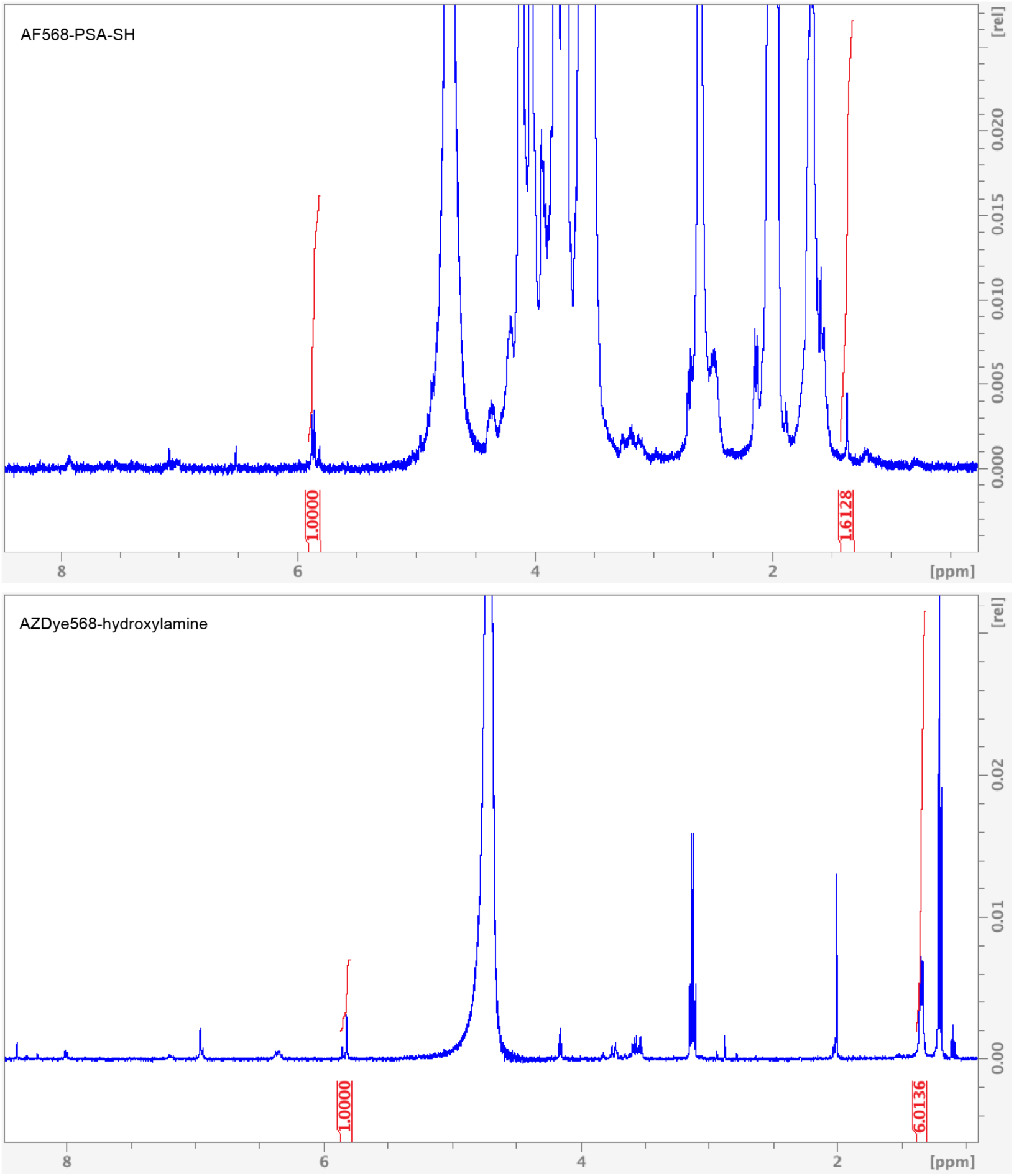
^1^H NMR analysis of 568-PSA-Tet and AZDye568-hydroxylamine. ^1^H NMR signals δ5.8 ppm (aromatic and alkene protons) represent pendant AZDye568 in D_2_O.

**Figure S19.**
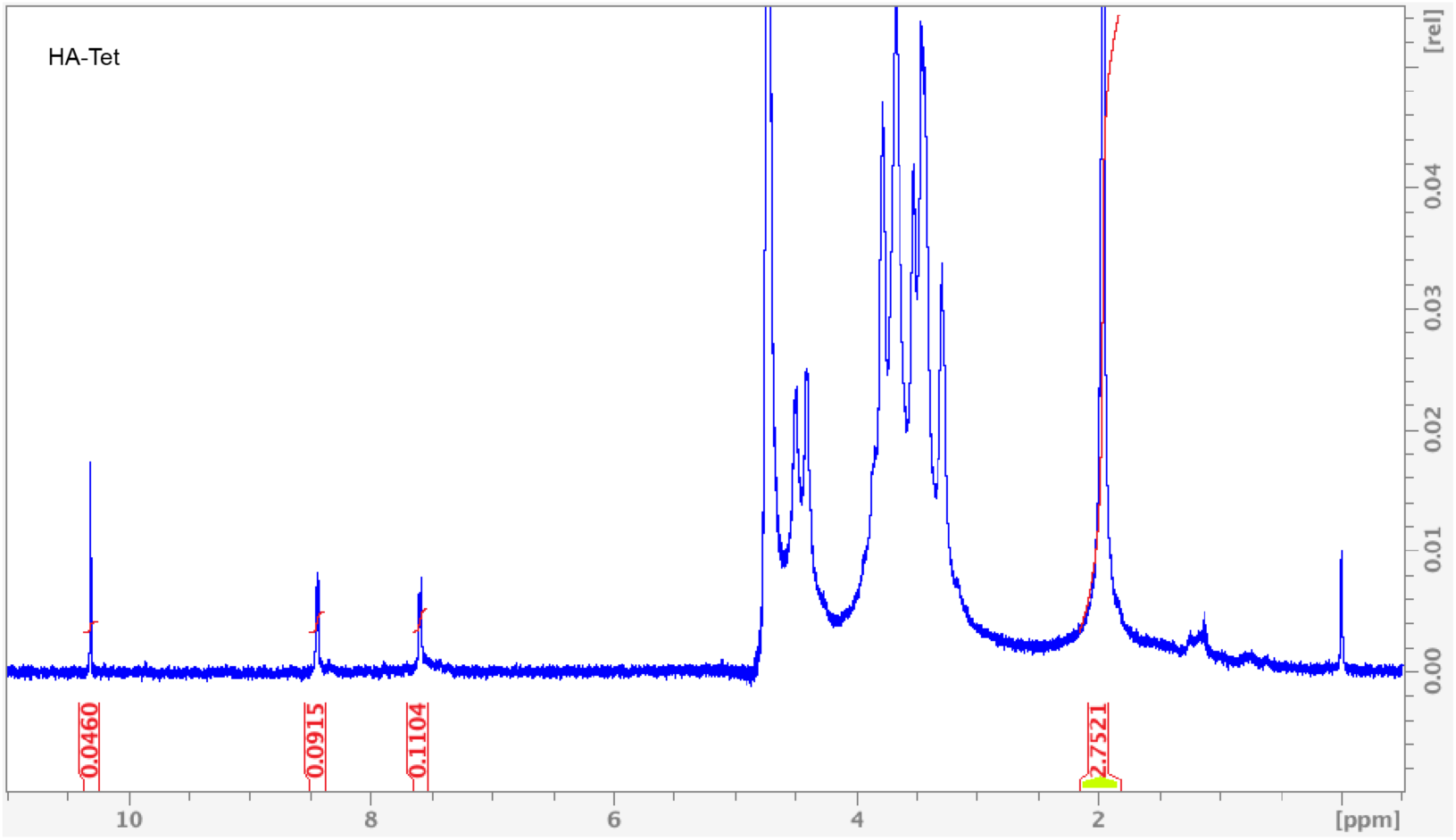
^1^H NMR analysis of HA-Tet. ^1^H NMR signals at δ8.5 (2H) and δ7.7 (2H) (aromatic protons) represent tetrazines in D_2_O.

**Figure 20.**
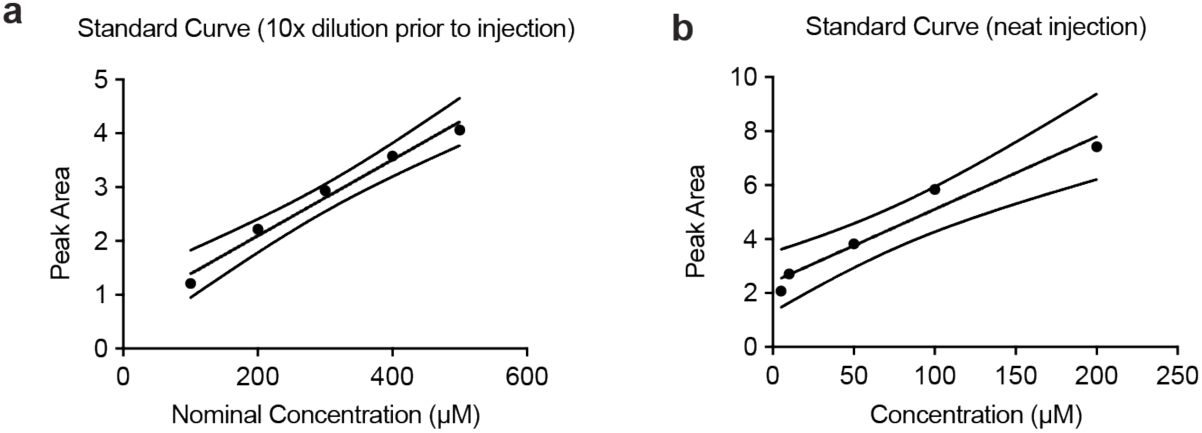
HPLC Standard Curves for Quantification of Free Neu5Ac. **a**, Standard curve generated from HPLC analysis of known Neu5Ac concentrations with a 10-fold dilution prior to injection, utilized for quantifying released Neu5Ac from soluble PSA polymer degradation assays. **b**, Standard curve from undiluted (neat) Neu5Ac standards, utilized for precise quantification of Neu5Ac release from MAP scaffold supernatant. Data points represent individual measurements plotted with linear regression fits (R² values shown).

## Reference

[1] a)K. W. Moremen, M. Tiemeyer, A. V. Nairn, Nature Reviews Molecular Cell Biology 2012, 13, 448; b)C. Reily, T. J. Stewart, M. B. Renfrow, J. Novak, *Nature Reviews Nephrology* 2019, 15, 346; c)A. Varki, *Glycobiology* 2016, 27, 3; d)V. Sytnyk, I. Leshchyns’ka, M. Schachner, *Cell Mol Life Sci* 2021, 78, 93; e)H. H. Freeze, E. A. Eklund, B. G. Ng, M. C. Patterson, *Lancet Neurol* 2012, 11, 453; f)R. Kleene, M. Schachner, *Nat Rev Neurosci* 2004, 5, 195.

[2] a)L. Silpa, R. Sim, A. J. Russell, Bioorganic & Medicinal Chemistry Letters 2022, 61, 128601; b)J. Wang, S. Chen, C. Pan, G. Li, Z. Tang, *Front Bioeng Biotechnol* 2022, 10, 799152; c)S. Jarrin, S. Cabré, E. Dowd, *Neurochemistry International* 2021, 144, 104971; d)Z. Giorgi, V. Veneruso, E. Petillo, P. Veglianese, G. Perale, F. Rossi, *ACS Applied Bio Materials* 2024, 7, 80; e)I. A. Hammam, R. Winters, Z. Hong, *Biomedical Engineering Advances* 2024, 8, 100132; f)G. Potjewyd, S. Moxon, T. Wang, M. Domingos, N. M. Hooper, *Trends Biotechnol* 2018, 36, 457.

[3] a)Z. Giorgi, V. Veneruso, E. Petillo, P. Veglianese, G. Perale, F. Rossi, ACS Appl Bio Mater 2024, 7, 80; b)A. Guijarro-Belmar, A. Varone, M. R. Baltzer, S. Kataria, E. Tanriver- Ayder, R. Watzlawick, E. Sena, C. J. Cunningham, A. M. Rajnicek, M. Macleod, W. Huang, G. L. Currie, S. K. McCann, *Spinal Cord* 2022, 60, 1041.

[4] a)P. R. Crocker, J. C. Paulson, A. Varki, Nature Reviews Immunology 2007, 7, 255; b)S. Duan, J. C. Paulson, *Annu Rev Immunol* 2020, 38, 365; c)V. S. Mahajan, S. Pillai, *Immunol Rev* 2016, 269, 145.

[5] a)A. Krishnan, V. G. Sendra, D. Patel, A. Lad, M. K. Greene, P. Smyth, S. A. Gallaher, M. Herron Ú, C. J. Scott, M. Genead, M. Tolentino, Front Immunol 2023, 14, 1237016; b)S. L. Peterson, A. Krishnan, D. Patel, A. Khanehzar, A. Lad, J. Shaughnessy, S. Ram, D. Callanan, D. Kunimoto, M. A. Genead, M. J. Tolentino, *Pharmaceuticals (Basel)* 2024, 17; c)X.-J. Wang, C.-H. Peng, S. Zhang, X.-L. Xu, G.-F. Shu, J. Qi, Y.-F. Zhu, D.-M. Xu, X.-Q. Kang, K.-J. Lu, F.-Y. Jin, R.-S. Yu, X.-Y. Ying, J. You, Y.-Z. Du, J.-S. Ji, *Nano Letters* 2019, 19, 829; d)S. Zhang, X. J. Wang, W. S. Li, X. L. Xu, J. B. Hu, X. Q. Kang, J. Qi, X. Y. Ying, J. You, Y. Z. Du, *Acta Biomater* 2018, 77, 15.

[6] Y. Su, C. Guo, Q. Chen, H. Guo, J. Wang, M. Kaihang, D. Chen, Int J Biol Macromol 2022, 214, 278.

[7] M. Mühlenhoff, M. Rollenhagen, S. Werneburg, R. Gerardy-Schahn, H. Hildebrandt, Neurochemical Research 2013, 38, 1134.

[8] a)L. R. Nih, E. Sideris, S. T. Carmichael, T. Segura, Advanced Materials 2017, 29, 1606471; b)K. Erning, K. L. Wilson, C. S. Smith, L. Nguyen, N. I. Joseph, R. Irengo, L. Y. Cao, M. Cumaran, Y. Shi, S. Lyu, L. Riley, T. W. Dunn, S. T. Carmichael, T. Segura, *Advanced Functional Materials*, n/a, 2500696.

[9] a)L. R. Conroy, T. R. Hawkinson, L. E. A. Young, M. S. Gentry, R. C. Sun, Trends Endocrinol Metab 2021, 32, 980; b)S. W. Yoo, M. G. Motari, K. Susuki, J. Prendergast, A. Mountney, A. Hurtado, R. L. Schnaar, *Faseb j* 2015, 29, 3040; c)R. L. Schnaar, R. Gerardy-Schahn, H. Hildebrandt, *Physiol Rev* 2014, 94, 461; d)A. V. Pshezhetsky, L. I. Ashmarina, *Biochemistry (Mosc)* 2013, 78, 736.

[10] J. Tena, C. B. Lebrilla, Proc Natl Acad Sci U S A 2021, 118.

[11] a)M. Noel, S. Suttapitugsakul, R. D. Cummings, R. G. Mealer, Proceedings of the National Academy of Sciences 2025, 122, e2418949122; b)Q. Zhang, C. Ma, L. S. Chin, S. Pan, L. Li, *Sci Adv* 2024, 10, eadk6911; c)J. Tena, I. Maezawa, M. Barboza, M. Wong, C. Zhu, M. R. Alvarez, L.-W. Jin, A. M. Zivkovic, C. B. Lebrilla, *Molecular & Cellular Proteomics* 2022, 21; d)S. Zamze, D. J. Harvey, Y. J. Chen, G. R. Guile, R. A. Dwek, D. R. Wing, *Eur J Biochem* 1998, 258, 243; e)S. E. Williams, M. Noel, S. Lehoux, M. Cetinbas, R. J. Xavier, R. I. Sadreyev, E. M. Scolnick, J. W. Smoller, R. D. Cummings, R. G. Mealer, *Nat Commun* 2022, 13, 275; f)S. E. Williams, M. Noel, S. Lehoux, M. Cetinbas, R. J. Xavier, R. I. Sadreyev, E. M. Scolnick, J. W. Smoller, R. D. Cummings, R. G. Mealer, *Nature Communications* 2022, 13, 275; g)R. Beatson, V. Tajadura-Ortega, D. Achkova, G. Picco, T. D. Tsourouktsoglou, S. Klausing, M. Hillier, J. Maher, T. Noll, P. R. Crocker, J. Taylor-Papadimitriou, J. M. Burchell, *Nat Immunol* 2016, 17, 1273; h)Y. Pan, T. Yago, J. Fu, B. Herzog, J. M. McDaniel, P. Mehta-D’Souza, X. Cai, C. Ruan, R. P. McEver, C. West, K. Dai, H. Chen, L. Xia, *Blood* 2014, 124, 3656.

[12] a)J. V. Pluvinage, M. S. Haney, B. A. H. Smith, J. Sun, T. Iram, L. Bonanno, L. Li, D. P. Lee, D. W. Morgens, A. C. Yang, S. R. Shuken, D. Gate, M. Scott, P. Khatri, J. Luo, C. R. Bertozzi, M. C. Bassik, T. Wyss-Coray, Nature 2019, 568, 187; b)T. Fujioka, N. Kaneko, I. Ajioka, K. Nakaguchi, T. Omata, H. Ohba, R. Fässler, J. M. García-Verdugo, K. Sekiguchi, N. Matsukawa, K. Sawamoto, *EBioMedicine* 2017, 16, 195.

[13] W. Zhu, Y. Zhou, L. Guo, S. Feng, Cell Death Discovery 2024, 10, 415.

[14] a)D. H. Allendorf, E. H. Franssen, G. C. Brown, Journal of Neurochemistry 2020, 155, 403; b)S. Zeng, Y. Wen, C. Yu, *Cellular Signalling* 2023, 112, 110927; c)L. Nanetti, A. Vignini, F. Raffaelli, R. Taffi, M. Silvestrini, L. Provinciali, L. Mazzanti, *Dis Markers* 2008, 25, 167.

[15] N. Mukherjee, S. Nandi, S. Garg, S. Ghosh, S. Ghosh, R. Samat, S. Ghosh, ACS Chemical Neuroscience 2020, 11, 231.

[16] a)T. M. Villanueva-Cabello, L. D. Gutiérrez-Valenzuela, R. Salinas-Marín, D. V. López-Guerrero, I. Martínez-Duncker, Frontiers in Immunology 2022, Volume 12 - 2021; b)H. Thiesler, H. Hildebrandt, Neural Regeneration Research 2026, 21, 661; c)L.-J. Schröder, H. Thiesler, L. Gretenkort, T. M. Möllenkamp, M. Stangel, V. Gudi, H. Hildebrandt, *Frontiers in Cellular Neuroscience* 2023, Volume 17 - 2023.

[17] a)L.-J. Kang, E. Oh, C. Cho, H. Kwon, C.-G. Lee, J. Jeon, H. Lee, S. Choi, S. J. Han, J. Nam, C.-u. Song, H. Jung, H. Y. Kim, E.-J. Park, E.-J. Choi, J. Kim, S.-i. Eyun, S. Yang, Scientific Reports 2020, 10, 5603; b)M. B. Jones, *Cellular Immunology* 2018, 333, 58; c)Y. Ohmi, T. Nishikaze, Y. Kitaura, T. Ito, S. Yamamoto, F. Sugiyama, M. Matsuyama, Y. Takahashi, A. Takeda, T. Kawahara, T. Okajima, K. Furukawa, K. Furukawa, *Glycobiology* 2021, 31, 557.

[18] S. Grabenstein, K. N. Barnard, M. Anim, A. Armoo, W. S. Weichert, C. R. Bertozzi, C. R. Parrish, R. Willand-Charnley, Glycobiology 2021, 31, 1279.

[19] a)T. Paramo, T. J. Piggot, C. E. Bryant, P. J. Bond, J Biol Chem 2013, 288, 36215; b)C. O. Soares, A. S. Grosso, J. Ereño-Orbea, H. Coelho, F. Marcelo, *Front Mol Biosci* 2021, 8, 727847; c)J. R. Caso, J. M. Pradillo, O. Hurtado, P. Lorenzo, M. A. Moro, I. Lizasoain, *Circulation* 2007, 115, 1599; d)G. Y. Chen, N. K. Brown, W. Wu, Z. Khedri, H. Yu, X. Chen, D. van de Vlekkert, A. D’Azzo, P. Zheng, Y. Liu, *Elife* 2014, 3, e04066.

[20] M. Teodorowicz, O. Perdijk, I. Verhoek, C. Govers, H. F. Savelkoul, Y. Tang, H. Wichers, K. Broersen, *PLoS One* 2017, 12, e0173778.

[21] C. Gauthier-Campbell, T. Lester, V. Sluzky, Pharmaceut Reg Affairs 2018, 7, 2.

[22] B. S. Park, J.-O. Lee, *Experimental & Molecular Medicine* 2013, 45, e66.

[23] a)K. Boelaars, Y. van Kooyk, Trends in Cancer 2024, 10, 230; b)R. Rashmi, B. P. Bode, N. Panesar, S. B. King, J. R. Rudloff, M. R. Gartner, J. M. Koenig, *Pediatric Research* 2009, 66, 266; c)J. Q. Zhang, B. Biedermann, L. Nitschke, P. R. Crocker, *Eur J Immunol* 2004, 34, 1175.

[24] a)Y. Wu, D. Yang, R. Liu, L. Wang, G.-Y. Chen, iScience 2020, 23; b)T. J. Borges, K. Lima, R. B. Gassen, K. Liu, Y. Ganchiku, G. T. Ribas, M. Liao, J. I. B. Goncalves, I. T. Lape, I. A. Rosales, Y. Zhao, E. Hui, R. L. Fairchild, C. LeGuern, C. Bonorino, S. K. Calderwood, J. C. Madsen, L. V. Riella, Science Translational Medicine 2025, 17, eads2694; c)C. R. Boyd, S. J. Orr, S. Spence, J. F. Burrows, J. Elliott, H. P. Carroll, K. Brennan, N. G. J, W. A. Coulter, C. Jones, P. R. Crocker, J. A. Johnston, C. A. Jefferies, *J Immunol* 2009, 183, 7703.

[25] a)N. F. EFSA Panel on Nutrition, F. Allergens, D. Turck, J. Castenmiller, S. De Henauw, K. I. Hirsch-Ernst, J. Kearney, A. Maciuk, I. Mangelsdorf, H. J. McArdle, A. Naska, C. Pelaez, K. Pentieva, A. Siani, F. Thies, S. Tsabouri, M. Vinceti, F. Cubadda, K. H. Engel, T. Frenzel, M. Heinonen, R. Marchelli, M. Neuhäuser-Berthold, M. Poulsen, J. R. Schlatter, H. van Loveren, P. Colombo, H. K. Knutsen, EFSA Journal 2020, 18, e06098; b)A. E. Manzi, H. H. Higa, S. Diaz, A. Varki, *J Biol Chem* 1994, 269, 23617.

[26] K. R. Reiding, D. Blank, D. M. Kuijper, A. M. Deelder, M. Wuhrer, Anal Chem 2014, 86, 5784.

[27] a)D. Liu, Z. Zhao, A. Wang, S. Ge, H. Wang, X. Zhang, Q. Sun, W. Cao, M. Sun, L. Wu, M. Song, Y. Zhou, W. Wang, Y. Wang, Journal of Neuroinflammation 2018, 15, 123; b)J. Li, Y. Qiu, C. Zhang, H. Wang, R. Bi, Y. Wei, Y. Li, B. Hu, *Pharmacological Research* 2023, 191, 106726.

[28] M. Y. Lee, J. A. Yang, H. S. Jung, S. Beack, J. E. Choi, W. Hur, H. Koo, K. Kim, S. K. Yoon, S. K. Hahn, ACS Nano 2012, 6, 9522.

[29] Y. Zeng, T. N. Ramya, A. Dirksen, P. E. Dawson, J. C. Paulson, Nat Methods 2009, 6, 207.

[30] S. D. Fontaine, R. Reid, L. Robinson, G. W. Ashley, D. V. Santi, Bioconjugate Chemistry 2015, 26, 145.

[31] a)K. Zlatina, M. Saftenberger, A. Kühnle, C. E. Galuska, U. Gärtner, A. Rebl, M. Oster, A. Vernunft, S. P. Galuska, International Journal of Molecular Sciences 2018, 19, 1679; b)A. Shahraz, Y. Lin, J. Mbroh, J. Winkler, H. Liao, M. Lackmann, A. Bungartz, P. F. Zipfel, C. Skerka, H. Neumann, *Scientific Reports* 2022, 12, 5818; c)I. Oltmann-Norden, S. P. Galuska, H. Hildebrandt, R. Geyer, R. Gerardy-Schahn, H. Geyer, M. Mühlenhoff, *J Biol Chem* 2008, 283, 1463; d)C. Sato, H. Fukuoka, K. Ohta, T. Matsuda, R. Koshino, K. Kobayashi, F. A. Troy, K. Kitajima, *Journal of Biological Chemistry* 2000, 275, 15422.

[32] K. L. Wilson, S. C. L. Pérez, M. M. Naffaa, S. H. Kelly, T. Segura, Advanced Materials 2022, 34, 2201921.

[33] a)J. Exton, J. M. G. Higgins, J. Chen, Scientific Reports 2023, 13, 12826; b)P. C. Georges, W. J. Miller, D. F. Meaney, E. S. Sawyer, P. A. Janmey, *Biophys J* 2006, 90, 3012.

[34] X. Zhao, N. Huebsch, D. J. Mooney, Z. Suo, J Appl Phys 2010, 107, 63509.

[35] L. Riley, P. Cheng, T. Segura, Nature Computational Science 2023, 3, 975.

[36] Y. Liu, A. Suarez-Arnedo, L. Riley, T. Miley, J. Xia, T. Segura, Adv Healthc Mater 2023, 12, e2300823.

[37] L. Nanetti, A. Vignini, F. Raffaelli, R. Taffi, M. Silvestrini, L. Provinciali, L. Mazzanti, Disease Markers 2008, 25, 613272.

[38] E. A. Goode, M. Orozco-Moreno, K. Hodgson, A. Nabilah, M. Murali, Z. Peng, J. Merx, E. Rossing, J. F. A. Pijnenborg, T. J. Boltje, N. Wang, D. J. Elliott, J. Munkley, Cancers (Basel) 2024, 16.

[39] A. Shahraz, J. Kopatz, R. Mathy, J. Kappler, D. Winter, S. Kapoor, V. Schütza, T. Scheper, V. Gieselmann, H. Neumann, Scientific Reports 2015, 5, 16800.

[40] a)E. Natkanski, W. Y. Lee, B. Mistry, A. Casal, J. E. Molloy, P. Tolar, Science 2013, 340, 1587; b)L. X. Wang, S. X. Zhang, H. J. Wu, X. L. Rong, J. Guo, *J Leukoc Biol* 2019, 106, 345; c)Y. Yue, S. Huang, H. Li, W. Li, J. Hou, L. Luo, Q. Liu, C. Wang, S. Yang, L. Lv, J. Shao, Z. Wu, *Annals of Translational Medicine* 2020, 8, 1409; d)Z. Strizova, I. Benesova, R. Bartolini, R. Novysedlak, E. Cecrdlova, Lily K. Foley, I. Striz, *Clinical Science* 2023, 137, 1067; e)M. S. Macauley, P. R. Crocker, J. C. Paulson, *Nature Reviews Immunology* 2014, 14, 653.

[41] L. Li, Y. Chen, M. N. Sluter, R. Hou, J. Hao, Y. Wu, G.-Y. Chen, Y. Yu, J. Jiang, Journal of Neuroinflammation 2022, 19, 191.

[42] L. Liu, J. V. Stokes, W. Tan, S. B. Pruett, Journal of Immunological Methods 2022, 511, 113378.

[43] N. V. Phan, E. M. Rathbun, Y. Ouyang, S. T. Carmichael, T. Segura, Nature Reviews Bioengineering 2024, 2, 44.

[44] G. C. Jickling, D. Liu, B. P. Ander, B. Stamova, X. Zhan, F. R. Sharp, J Cereb Blood Flow Metab 2015, 35, 888.

[45] a)Y. Wang, R. K. Leak, G. Cao, Front Cell Neurosci 2022, 16, 980722; b)J. E. Anttila, K. W. Whitaker, E. S. Wires, B. K. Harvey, M. Airavaara, *Prog Neuropsychopharmacol Biol Psychiatry* 2017, 79, 3; c)J. Jia, L. Zheng, L. Ye, J. Chen, S. Shu, S. Xu, X. Bao, S. Xia, R. Liu, Y. Xu, M. Zhang, *Cell Death & Disease* 2023, 14, 156; d)W. Zhang, J. Zhao, R. Wang, M. Jiang, Q. Ye, A. D. Smith, J. Chen, Y. Shi, *CNS Neurosci Ther* 2019, 25, 1329; e)T. Shibahara, T. Ago, M. Tachibana, K. Nakamura, K. Yamanaka, J. Kuroda, Y. Wakisaka, T. Kitazono, *Stroke* 2020, 51, 3095.

[46] C. Chen, T. Huang, X. Zhai, Y. Ma, L. Xie, B. Lu, Y. Zhang, Y. Li, Z. Chen, J. Yin, P. Li, J Cereb Blood Flow Metab 2021, 41, 2150.

[47] a)T. Li, T. Xu, J. Zhao, H. Gao, W. Xie, Free Radical Biology and Medicine 2022, 181, 209; b)J. M. Wierońska, P. Cieślik, L. Kalinowski, *Biomolecules* 2021, 11, 1097.

[48] a)T. R. Sippel, T. Shimizu, F. Strnad, R. J. Traystman, P. S. Herson, A. Waziri, Journal of Cerebral Blood Flow & Metabolism 2015, 35, 1657; b)A. B. Petrone, G. C. O’Connell, M. D. Regier, P. D. Chantler, J. W. Simpkins, T. L. Barr, *Transl Stroke Res* 2016, 7, 103.

[49] a)T. Li, J. Zhao, H. Gao, Int J Mol Sci 2022, 23; b)Y. Wang, R. K. Leak, G. Cao, *Frontiers in Cellular Neuroscience* 2022, Volume 16 - 2022.

[50] a)X. Hu, R. K. Leak, Y. Shi, J. Suenaga, Y. Gao, P. Zheng, J. Chen, Nat Rev Neurol 2015, 11, 56; b)R. M. Ritzel, A. R. Patel, J. M. Grenier, J. Crapser, R. Verma, E. R. Jellison, L. D. McCullough, *J Neuroinflammation* 2015, 12, 106; c)E. E. Wicks, K. R. Ran, J. E. Kim, R. Xu, R. P. Lee, C. M. Jackson, *Frontiers in Immunology* 2022, Volume 13 - 2022.

[51] S. W. Hsieh, L. C. Huang, Y. P. Chang, C. H. Hung, Y. H. Yang, Psychiatry Clin Neurosci 2020, 74, 383.

[52] T. R. Sippel, T. Shimizu, F. Strnad, R. J. Traystman, P. S. Herson, A. Waziri, J Cereb Blood Flow Metab 2015, 35, 1657.

[53] Y. Liu, A. Suarez-Arnedo, L. Riley, T. Miley, J. Xia, T. Segura, Advanced Healthcare Materials 2023, 12, 2300823.

[54] W. Ying, P. S. Cheruku, F. W. Bazer, S. H. Safe, B. Zhou, J Vis Exp 2013.

[55] C. E. Galuska, J. A. Dambon, A. Kühnle, K. F. Bornhöfft, G. Prem, K. Zlatina, T. Lütteke, S. P. Galuska, Front Immunol 2017, 8, 1229.

[56] a)A. Agarwal, B. G. Bobay, M. L. Becker, Journal of the American Chemical Society 2025, 147, 9386; b)W. Li, H. Chung, C. Daeffler, J. A. Johnson, R. H. Grubbs, *Macromolecules* 2012, 45, 9595.

[57] C. Stringer, T. Wang, M. Michaelos, M. Pachitariu, Nature Methods 2021, 18, 100.

[58] A. R. Anderson, E. Nicklow, T. Segura, Acta Biomaterialia 2022, 150, 111.

[59] A. R. Anderson, E. L. P. Caston, L. Riley, L. Nguyen, D. Ntekoumes, S. Gerecht, T. Segura, Advanced Functional Materials 2025, 35, 2400567.

